# IFDlong: an isoform and fusion detector for accurate annotation and quantification of long-read RNA-seq data

**DOI:** 10.1101/2024.05.11.593690

**Authors:** Wenjia Wang, Yuzhen Li, Sungjin Ko, Ning Feng, Manling Zhang, Jia-Jun Liu, Songyang Zheng, Baoguo Ren, Yan P. Yu, Jian-Hua Luo, George C. Tseng, Silvia Liu

## Abstract

Advancements in long-read transcriptome sequencing (long-RNA-seq) technology have revolutionized the study of isoform diversity. These full-length transcripts enhance the detection of various transcriptome structural variations, including novel isoforms, alternative splicing events, and fusion transcripts. By shifting the open reading frame or altering gene expressions, studies have proved that these transcript alterations can serve as crucial biomarkers for disease diagnosis and therapeutic targets. In this project, we proposed IFDlong, a bioinformatics and biostatistics tool to detect isoform and fusion transcripts using bulk or single-cell long-RNA-seq data. Specifically, the software performed gene and isoform annotation for each long-read, defined novel isoforms, quantified isoform expression by a novel expectation-maximization algorithm, and profiled the fusion transcripts. For evaluation, IFDlong pipeline achieved overall the best performance when compared with several existing tools in large-scale simulation studies. In both isoform and fusion transcript quantification, IFDlong is able to reach more than 0.8 Spearman’s correlation with the truth, and more than 0.9 cosine similarity when distinguishing multiple alternative splicing events. In novel isoform simulation, IFDlong can successfully balance the sensitivity (higher than 90%) and specificity (higher than 90%). Furthermore, IFDlong has proved its accuracy and robustness in diverse in-house and public datasets on healthy tissues, cell lines and multiple types of diseases. Besides bulk long-RNA-seq, IFDlong pipeline has proved its compatibility to single-cell long-RNA-seq data. This new software may hold promise for significant impact on long-read transcriptome analysis. The IFDlong software is available at https://github.com/wenjiaking/IFDlong.

## Introduction

Long-read technology has brought the high-throughput genomic sequencing into the third generation to study genomes, transcriptomes, and metagenomes. Long-read sequencing is able to generate reads up to several million base pairs, while short-read sequencing typically yields reads with only 50 to 300 base pairs. For transcriptome sequencing, long-read RNA-seq (long-RNA-seq) can successfully reveal the full-length transcripts at an unprecedented resolution. Compare with short-read RNA-seq, long-RNA-seq presents additional advantages. First, long-RNA-seq can reduce the noise caused by artificial amplification and can dramatically increase the alignment certainty. Second, by covering full-length (or almost full-length) of the transcript sequence, long reads not only enable the precise identification and quantification of known isoforms, but also show power to discover the novel isoforms. In addition, long reads have advantages to detect transcriptomic structural variants by precisely locating the exon donor-acceptor merging points for alternative splicing analyses or covering the fusion junction points for fusion transcript identification. Several platforms have facilitated this cutting-edge technique. Techniques from two companies have been widely applied for long-read sequencing: Pacific Biosciences (PacBio) Iso-Seq^1^ and Oxford Nanopore Technology (ONT) whole-transcriptome sequencing^2, 3^. Other than these direct long-read sequencing platforms, several synthetic long-read strategies have been developed as well, including 10X Genomics Linked-reads^4^ and Element Biosciences LoopSeq^5, 6^.

One of the major applications for long-RNA-seq is to analyze isoforms and transcriptomic structural variants (TSVs). TSV refers to the transcriptome sequence alterations that are caused by genomic structural variants (SVs) or transcriptomic splicing. It usually describes large variant with more than thousands of base pairs. Other than genomic-level analysis, transcriptome-level variant study can add flexibility to describe alternative splicing, differences in expression levels and potential functional alteration in protein expression. These variant events may play roles as biomarkers for disease diagnoses and serve as therapeutic targets. Two major TSVs will be explored in this study: alternative splicing and fusion transcript. Alternative splicing (or pre-mRNA splicing) refers to the exon splicing events on pre-mRNA, where exons can be alternatively included or removed when forming mature mRNA^7, 8^. Given the fact that over- or under-expressed isoforms (or proteins) and novel isoforms will influence the regulation system, deregulation of alternative splicing events is a hall mark of cancer. Thus, disease-associated isoform-specific events can serve as biomarkers for disease diagnosis and treatment^9-11^. For isoform study, limited by read length, short-read sequencing is not adequate to capture the full transcripts, which can only quantify the isoform abundance by statistical inference and hard to discover novel isoforms. In contrast, long read shows its advantage in identifying the exact isoform and cover the splicing points by full-length transcript. However, besides complete long-reads, truncated long-reads will be generated in the meanwhile, which will cause the uncertainty of isoform assignments (**Fig. 1A**). Thus, statistical estimation for isoform expression by long-RNA-seq is still urgently needed. Several bioinformatics tools have been developed to identify isoforms on long-RNA-seq data. LIQA^12^and Mandalorion^13^ are able to perform isoform quantification, but cannot perform isoform annotation for individual long-read. TALON^14^ can perform both, but its accuracy can be further improved. In addition, only TALON is able to discover novel isoforms, while LIQA and Mandalorion are merely focused on known ones.

**Figure 1:**
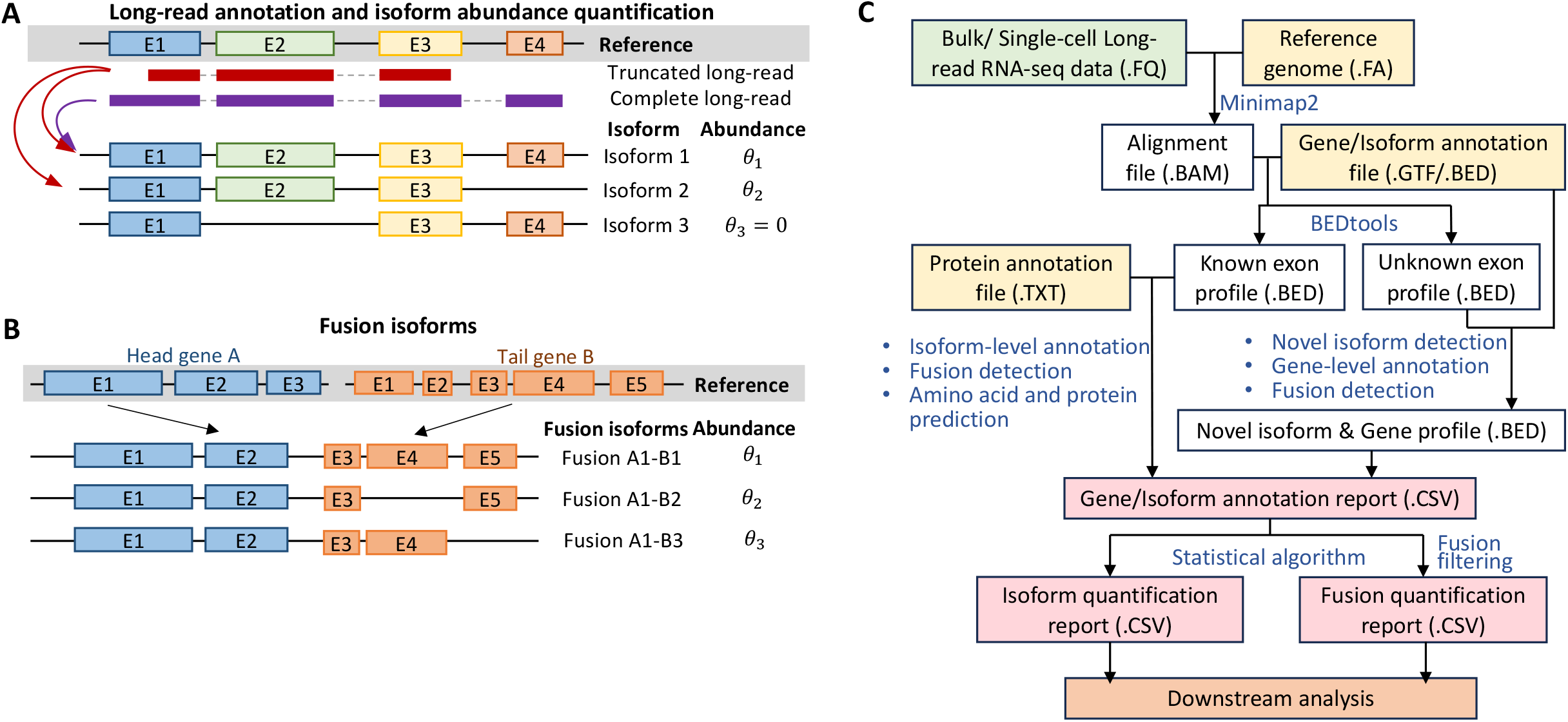
Illustration of alternative splicing, fusion isoform and IFDlong pipeline. (A) Long-read annotation and isoform abundance quantification. The red and purple reads represent the truncated and completed long-reads, which can serve as supporting reads by alignment to different isoforms. (B) Illustration of fusion isoforms. Head gene A and tail gene B will potentially form into different fusion isoforms. (C) IFDlong pipeline. The proposed pipeline will take in the raw long-read sequences and annotation files, and ultimately output the gene/isoform annotation reports, isoform quantification reports and fusion quantification reports for downstream analysis.

Fusion transcripts (**Fig. 1B**) are caused by fusion genes or trans-splicing events resulting from chromosomal rearrangements or ligation of two different primary RNA transcripts. Fusion transcripts play important roles in tumorigenesis by potentially shifting the open reading frame (ORF) for the tail protein, forming novel chimera proteins, or altering gene expression^15-18^. Previous studies have proved that fusions are highly associated with occurrence and recurrence of caners^19-21^. Suicide-gene insertion on the fusion gene can serve as genotype-specific cancer therapy. In addition to the DNA-level of fusion gene study, fusion transcript analysis can reveal multiple alternative splicing events^22^ As the toy example shown in **Fig. 1B**, gene A can fuse with three different isoforms of gene B. Short-read RNA-seq has been largely applied to discover fusions^23^. However, it has limitation in either performing isoform resolution analysis or requiring extremely deep sequencing (e.g. 1300x sequencing depth to detect rare fusion events in our previous study^24^). To overcome this, long reads possess advantage over short reads in capturing the fusion isoform and precisely locate the fusion junction point^25^. Currently, bioinformatics tools such as Genion^26^ and JAFFAL^27^, have been developed to call fusion transcripts from long-RNA-seq that. However, they can only detect fusion events at gene level, but not isoform resolution. Additionally, neither tool will perform fusion isoform quantification to estimate the expression level of normal and fusion isoforms. Tool such as AERON^28^ is newly developed and its Github script is still under development.

In this article, we proposed a new bioinformatics pipeline IFDlong, an <u>I</u>soform <u>F</u>usion <u>D</u>etector that was tailored for <u>long</u>-RNA-seq data for the annotation and quantification of isoform and fusion transcripts. As shown in **Fig. 1C**, the pipeline can take in either bulk or single-cell long-RNA-seq data. Based on the reference genome and gene/isoform annotation profile, the long reads were aligned to reference genome and were annotate to genes and isoforms. For the case where no known isoform/genes can be annotated to the long-read, novel isoforms were defined. Followed by statistical inference methods, isoform and fusion quantification profiles were generated as output. Compared with the existing tools, IFDlong presents four major advantages: (1) IFDlong is compatible with both bulk and single-cell long-RNA-seq data; (2) IFDlong is able to perform read-level gene/isoform annotation and detect novel isoforms; (3) IFDlong performs statistical estimation on the truncated long-reads with multiple alignment for high-accurate isoform and fusion quantification; (4) IFDlong is the only tool that can do discover fusion transcripts at isoform resolution. To test the performance of proposed pipeline, we first compared IFDlong with several existing tools in *in-silico* data generated by long-read simulator to evaluate their performance under different sequencing parameters. Next, multiple published long-read data sets were employed and proved that our tool is compatible with different long-read platforms and its ability to accurately profile alternative splicing and fusion events. In summary, IFDlong showed overall best performance in terms of isoform quantification, novel isoform detection, and fusion isoform quantification. IFDlong will serve as a generalizable tool for advanced long-RNA-seq data analysis.

## Methods and Materials

### Reference and annotation file preparation

IFDlong requires the reference genome in FASTA format, the transcript annotation profile in BED format, the gene annotation profile in BED format, the amino acid (AA) annotation profile in TXT format, pseudo gene database in RDS format, and the root gene symbols of all gene families in TXT format. All the reference and annotation files for human and mouse are well prepared (available for download in Github) and will be fully described below. If the users are working on species other than human and mouse, please apply the IFDlong refDataSetup.sh function to build up the reference files.

IFDlong can take in both raw sequencing read (in FASTQ format) and aligned file (in BAM format). If working on FASTQ file, a reference genome in FASTA format is required to call Minimap2 aligner ^29^ **(Fig. 1C)** For gene and isoform annotation, gene annotation profile in BED format is required by IFDlong. For example, the annotation file in GTF format can be downloaded from the UCSC genome reference Consortium (e.g. GRCh38 for human and GRCm38 for mouse downloaded from https://hgdownload2.soe.ucsc.edu/downloads.html). Then the GTF file will be formatted by IFDlong (via command IFDlong.sh) into BED format, where each row records the chromosome, start position, end position, name, score, and strand of each consisting exon. Ultimately, a known gene and a known transcript profile will be used by IFDlong pipeline for gene and transcript annotation. IFDlong will build the AA annotation profile per known transcripts. Based on the reference genome (in FASTA format, described above) and the transcript profile (in BED format, described above), IFDlong will first apply the *getfasta* function in *bedtools*^30^ to extract the DNA/RNA sequences per transcript, followed by the *translate* function in *Biostrings* R package to translate the DNA/RNA sequences into AA sequences. These AA sequences will be saved in the TXT format as the AA annotation profile to be used for the prediction of AA sequence of the long read.

To filter out false positive fusion candidates, database for gene families and pseudo genes are required by IFDlong. The human pseudo gene database was downloaded from Pseudogene.org (http://pseudogene.org/psicube/data/gencode.v10.pseudogene.txt), and the mouse pseudo gene database was collected from Mouse Genome Informatics (https://www.informatics.jax.org/downloads/reports/MGI_BioTypeConflict.rpt). To play a complementary role, additional pseudo genes were identified based on the gene descriptions obtained from *Bioconductor* packages *org.Hs.eg.db* (for human) and *org.Mm.eg.db* (for mouse). The pseudo gene names were saved in an RDS file. To collect the gene family information, human database was downloaded from the HGNC BioMart server (https://biomart.genenames.org), which contains the family names and common root gene symbols of each human gene family. For mouse gene family information, given no reliable open-source database was found to the best of our knowledge, the mouse genes were first mapped to human homologous genes by MGI Data and Statistical Reports (https://www.informatics.jax.org/downloads/reports/index.html). Then the mapped mouse genes will employ the same gene family database as the human one.

### Isoform quantity estimates

The IFDlong isoform quantification method is derived from the LIQA^12^, with improved estimation algorithm utilizing the results from the upstream isoform annotation analysis. Mathematically, given a gene of interest denoted by *g*, let *R*^*g*^ represent the set of reads (denoted by *r*) that are aligned to the gene *g*, and *I*^*g*^ is the set of known isoforms (denoted by *i*) from gene *g*. For a specific isoform *i ∈ I*^*g*^, let 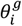 denote its relative abundance, with 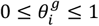 and 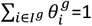. The probability that a read originates from isoform is defined as 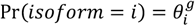 We further define the indicator matrix 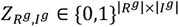 that is unobserved with entry *z*_*r,i*_*=* 1 if the read *r* is generated from a molecule that is originated from isoform *i*, and otherwise *z*_*r,i*_ *=* 0. For isoform quantification, the goal is to estimate the relative abundance 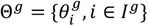 based on RNA-seq long reads annotated to the gene *g*.

With the notation above, the complete data likelihood of the long-RNA-seq aligned to gene *g* can be written as

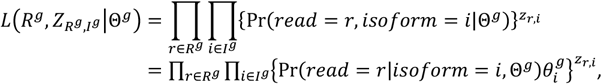

where 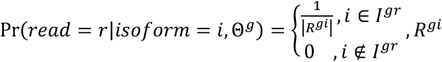 denotes the set of reads that are annotated to isoform *i* (i.e., a subset of *R*^*g*^, *R*^*gi*^ *⊂ R*^*g*^), and *I*^*g r*^ represents the set of isoforms that the read *r* is annotated to (similarly, *I*^*g r*^ *⊂ I*^*g*^). Note that it is possible that IFDlong annotates a read to multiple known isoforms which are highly overlapped with each other, and in this case, | *I*^*g r*^|>1.

Next, with the complete data likelihood, the update procedure of the Expectation-Maximization (EM) algorithm to estimate the parameter Θ^*g*^ is as follows:

### Initialization

Take the reads aligned to multiple isoforms as a supporting read for each of the isoforms (e.g., if one read is aligned to two isoforms of gene *g*, it will serve as a supporting read for both isoforms, as the red long-read illustrated in **Fig. 1A**), Θ^*g*^ will be initialized as proportion of the supporting reads aligned to each isoform for gene *g*.

### Expectation-step (E-step)

Calculate the expectation of the log likelihood with the following function:

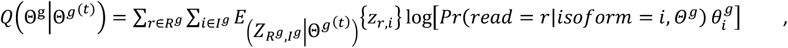

where 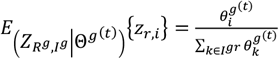

### Maximization-step (M-step)

Maximize function 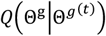 by Θ^*g*^ with the following formula,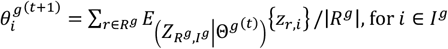.

The EM algorithm iteratively update the parameter by alternating between the E-step and M-step until convergence.

### Fusion quantity estimates

We use similar idea to perform fusion quantification. Given a gene fusion *f* consisting of all genes 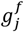 in gene set *G*^*f*^, let *R*^*f*^ denote the set of reads that are annotated to the gene fusion *f*, and *C*^*f*^ is the set of fusion transcripts from the gene fusion (i.e., each isoform component *i*_*cj*_ of a fusion transcript *c ∈ C*^*f*^ is from one of the gene 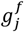 in *G*^*f*^). Then, we denote the relative abundance of each fusion transcript to be 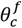 with 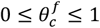 and 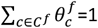, so the probability that a read originates from fusion transcript *t* is 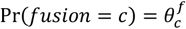. Similarly, we define the unobserved read-fusion transcript compatibility matrix 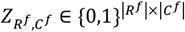 with entry *z*_*r,c*_ *=* 1 if the read *r* is generated from a molecule that is originated from fusion transcript *c*^*f*^, and otherwise *z*_*r,c*_ *=* 0. For fusion transcript quantification, our goal is to estimate the relative abundance 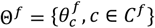 based on RNA-seq long reads annotated to the gene fusion *f*.

Similarly, the complete data likelihood is

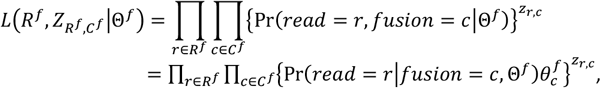

where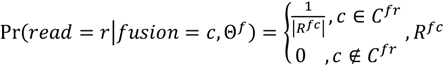, denotes the set of reads that are annotated to fusion transcript *c* (i.e., a subset of *R*^*f*^, *R*^*fc*^ *⊂ R*^*f*^), and *C*^*f r*^ represents the set of fusion transcripts that the read *r* is annotated to (similarly, *C*^*f r*^ *⊂ C*^*f*^). Again, it is possible that IFDlong annotates a read to multiple fusion transcripts which are are highly overlapped with each other, and in this case, |*C*^*f r*^|>1.

The EM algorithm iteratively update the parameter Θ^*f*^ by alternating between the E-step and M-step as below until convergence.

#### E-step

calculate the expectation of the log likelihood by

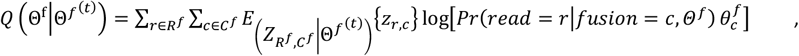

where 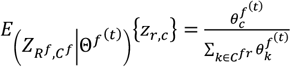

#### M-step

maximize function with resppect to Θ^*f*^ and have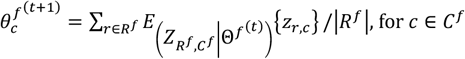

### Simulation data generation

In silico simulation data were generated to benchmark the software performance with underlying truth. Type I and Type F simulation datasets were generated for isoform and fusion evaluation, respectively.

The Type I1 simulation datasets were generated to mimic the distribution of whole transcriptome expression. First, to simulate the distribution of isoform expression in real data, Universal Human Reference (UHR) data was employed, and 28157 isoforms were detected and quantified (**Fig. S1A**). Second, sequence templates were constructed using these isoforms by synthesizing the exons of each isoform according to the transcript annotation profile. Next, multiple datasets of long-RNA-seq were generated by long-read simulator PBSIM2^31^ with different accuracy settings and gradient mean read lengths. Three accuracy setting were generated by adjusting the PBSIM2 parameters: accurate sequencing (*--difference-ratio=1:1:0* for substitution:insertion:deletion, and *--accuracy-mean=0.99*); ONT-like accuracy (*--difference-ratio=23:31:46* and *--accuracy-mean=0.85)*; and PacBio-like accuracy (*--difference-ratio=6:50:54* and *--accuracy-mean=0.85)*. Under the setting of high-accurate sequencing, simulation datasets with gradient mean read lengths were generated by setting *--length-mean* to be 200bps, 500bps, 800bps, 1000bps, 1200bps, 1500bps, 1800bps, and 2000bps (**Fig. S2**). Finally, each pool of simulated reads with a specific accuracy and mean read length setting was subsampled to gradient sequencing depth with 0.5 million, 1 million, 2 million and 4 million reads, respectively (**Fig. S1B**). Note that, the expression intensities of the 28157 isoforms follow the distribution of isoform expressions in the UHR data. In the meanwhile, the transcript and gene origin of each simulated long read was recorded to serve as the underlying truth.

The Type I2 simulation datasets were designed to generate long reads from multiple isoforms that were sourcing from the same gene. The sequence template consists of 16 transcripts of gene CTCFL (NM_001269043, NM_001269040, NM_001269041, NM_001269042, NM_080618, NM_001269046, NM_001269044, NM_001269049, NM_001269054, NM_001269045, NM_001269050, NM_001269048, NM_001269047, NM_001269055, NM_001269051, NM_001269052). Next, similar as the Type I1 simulation, multiple Type I2 datasets were generated with sequencing depth of 400 under the gradient mean read lengths (200bps, 500bps, 800bps, 1000bps, 1200bps, 1500bps, 1800bps, and 2000bps) and the three accuracy settings (accurate, PacBio and ONT).

The Type I3 simulation datasets aim to benchmark the novel isoform detection. First, the sequence templates were constructed by two isoform sets: 1000 known isoforms and novel isoforms. Novel isoforms were derived from the 1000 known transcripts with six different types: 859 templates with skipping exon, 1406 templates with additional exon, 873 templates with truncated alternative 5’ splice site, 1329 templates with extended alternative 5’ splice site, 873 templates with truncated alternative 3’ splice site, and 1325 templates with extended alternative 3’ splice site. Each individual type of novel isoforms was pooled with the 1000 known transcripts to generate six sequence templates in total. Per novelty type, simulator PBSIM2 was employed to generate long-RNA-seq datasets with sequencing depth of 40 under the mean read length of 1000bps and 2000bps, and the three different accuracy settings. The origin source and the novel type of each simulated long read were recorded to serve as the truth for performance evaluation.

The Type F1 datasets were simulated for fusion transcript quantification. The sequence templates consist of two sets of isoforms: 1000 selected known isoforms sourcing from different genes, and 1000 mosaic fusion transcript fusing two known isoforms from the 1000 known list with random breakpoints. Based on these template sequences, PBSIM2 was employed to generate simulation datasets under three accuracy settings with mean read length of 1500bps and sequencing depth of 400. According to the simulation log files, the read fusion status (fusion transcripts or normal isoform) and the isoform/ gene origin information of each long-read sequence were collected as underlying truth.

The Type F2 simulation datasets were intentionally designed to mimic the genes fused with multiple mates. To simulate the gene pairs that have high occurrence in real data, the TCGA fusion database (https://www.kobic.re.kr/chimerdb/download) was used as reference. Among the top 10 cancer types with the highest number of fusion transcripts, top 5 genes (referred as popular genes) with the highest number of paired genes were selected: FBXL20 (Breast Invasive Carcinoma, BRCA), CDS1 (Bladder Urothelial Carcinoma, BLCA), TMPRSS2 (Prostate Adenocarcinoma, PRAD), FRS2 (Sarcoma, SARC) and SFTPB (Lung Adenocarcinoma, LUAD). All the mate genes for these five popular genes are listed in the **Table S7**. For each popular gene, the fusion templates were constructed by fusing the popular gene and their mates according to the fusion breakpoints in the TCGA fusion database. Then based on these fusion templates, long-RNA-seq datasets were simulated under different mean read lengths (200bps, 1000bps, and 1500bps) and three accuracy settings (accurate, PacBio and ONT) with sequencing depth of 40 by PBSIM2. In the meanwhile, the fusion origins and breakpoint of each simulated long-read sequence was collected as the truth.

### Real data collection and pre-processing

#### Universal Human Reference (UHR)

Long-RNA-seq on the UHR sample was used to evaluate isoform quantification. The direct mRNA data sequenced by ONT platform was download from Gene Expression Omnibus (GEO) PRJNA639366^12^ with in total 476,000 reads and 896 base pairs in median, and 441,138 reads were aligned to human reference by Minimap2 aligner^29^. The UHR RNA (Agilent) + SIRV Isoform Mix E0 (Lexogen) sample measured by PacBio platform was downloaded from the PacBio website (https://downloads.pacbcloud.com/public/dataset/UHR_IsoSeq/). In total 6,775,127 reads with 1835 base pairs in median length were collected, and 6,385,883 reads was aligned to human reference by Minimap2 aligner. To serve as the underlying truth, measurements by the TaqMan Real-Time PCR Assays on the Stratagene UHR RNA samples were collected from the MicroArray Quality Control (MAQC) project via GEO GSE5350^32^. The geometric mean of the four repeating measurements was calculated as the true expression. In total 1044 isoforms were quantified and 964 of them have passed detection flag that will be used for this project.

#### Mouse data

For isoform quantification, long-RNA-seq data on mouse C2C12 cell line and heart tissue was downloaded from ENCODE project^33^. For C2C12, four repeats were downloaded from GEO database with accession ID GSE219813 and GSE219595. For mouse heart data, two repeats were downloaded from GSE219825. The isoform expression of both mouse datasets were quantified by TALON pipeline^14^.

#### Single-cell long-RNA-seq on hepatocellular carcinoma (HCC) patient

The data was downloaded from GEO GSE223743, including a paired liver benign – tumor samples from an HCC patient^34^. This dataset was applied for isoform quantification, differential expression analysis and fusion detection analysis. Single cells with at least 1000 long-reads were defined as valid cells, and in total 162 and 285 cells were analyzed from benign and tumor library. For fusion transcript detection, three fusion transcripts (EML4 - ACTR2, CCDC127 - PDCD6, and FGG - PLG) were validated by Sanger sequencing in the previous study^34^ that will serve as the truth.

#### Colon cancer samples

Tissue samples were collected from 3 colon cancer patients, including benign colon tissues (N), primary colon cancer samples (T), and lymph node metastasis samples (M). The samples were sequenced by Element Biosciences LoopSeq platform and the long-read data was deposited to GEO database with accession GSE155921^6^. Among the 8 samples, in total 4 two-way fusions (STAMBPL1 - FAS, SMYD3 - ZNF124, VAPB - GNAS, and ECHDC1 - PTPRK) were detected and validated using Taqman qRT-PCR and Sanger sequencing^6^, which will serve as the underlying truth for this study.

#### MCF7 human breast cancer cell line

MCF7 RNA extraction sequenced by both PacBio and ONT platforms were downloaded for isoform and fusion analysis. For PacBio SMRT sequencing, long-read FASTQ files were downloaded from NCBI SRA database with accession ID SRP055913 (https://www.ncbi.nlm.nih.gov/sra/?term=SRP055913). The 113 runs were concatenated into one file for analysis. For SGNex ONT sequencing, data was downloaded from https://github.com/GoekeLab/sg-nex-data. MCF7-EV_directRNA, MCF7_cDNA, MCF7_directRNA, and MCF7_directcDNA with multiple replicates were used in this study. As the true isoform expression, the RNA-seq on MCF7 with RSEM quantification of transcripts^35^ downloaded from Cancer Cell Line Encyclopedia (CCLE)^36^ were employed. Both two-way and three-way fusions that have been summarized in the previous research were used as fusion truth^27^.

#### H838 human lung adenocarcinoma cell line

Single-cell long-RNA-seq data on H838 sequenced by ONT platform was downloaded from GEO database with accession ID GSE154869^37^. Both two-way and three-way fusion transcripts were detected from this dataset for performance evaluation.

### Performance evaluation

#### Spearman’s correlation

is a nonparametric measure of statistical dependence between the rankings of two vectors. The correlation ranges from -1 to 1, with 1 indicting the same ranking between the two vectors, and -1 representing a fully opposed correlation. For both isoform or fusion transcript quantification, the Spearman’s correlation between the true number of supporting reads (denoted as vector 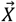) and the detected number of supporting reads (denoted as vector 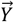) by each algorithm can be evaluated by the formula:

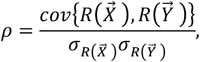

where 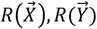 are the ranks of 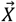 and 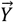 respectively; 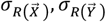 are the standard deviations of the rank variables, and 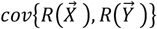 is the covariance of the rank variables.

#### Root Mean Squared Error (RMSE)

is employed to measure the differences between true and estimated values. Following the above notation, the RMSE is applied to benchmark the isoform quantification in the Type I1 simulation data by the formula:

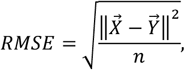

where *‖ · ‖* is *l*_2_-norm operator, and *n* is the length of the vector 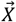 and 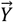.

#### Consistency measurement

*−log*10(*ave rage RMSE*) is used to evaluate the agreement across multiple repeats for isoform quantification. Isoform proportions were first averaged across all the repeats. Then for each repeat, the root mean squared error (RMSE) of the estimated isoform proportions was calculated compared to this mean proportion. Ultimately, RMSE values of all the repeats were averaged and -log10 transformed as described below,

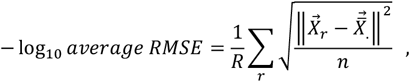

where *‖ · ‖* is *l*_2_ -norm operator; *n* is the length of the isoform proportion vector 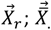 is the mean of 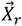 over *r =* {1,2, …, *R*}, and *R* is the number of repeats (e.g. four repeats for C2C12 mouse cell line or two repeats for mouse heart dataset).

#### Sensitivity and Specificity

Sensitivity (true positive rate) is the probability of a positive test result, conditioned on the individual truly being positive. Specificity (true negative rate) is the probability of a negative test result, conditioned on the individual truly being negative. To benchmark novel isoforms detection in the Type I3 simulation data, the sensitivity and specificity are calculated as:

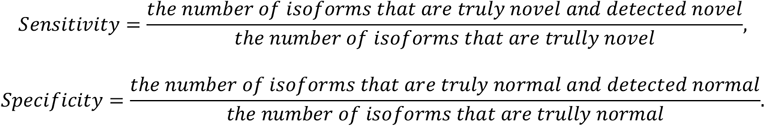

#### Precision, Recall and F1 score

Precision (also called positive predictive value) is the fraction of relevant instances among the retrieved instances. Recall (also known as sensitivity) is the fraction of relevant instances that were retrieved. The precision and recall are defined as:

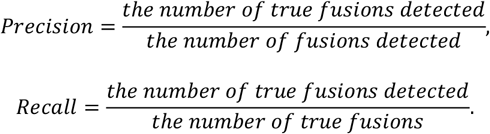

To balance the precision and recall, the F-measure (or F1 score) is the defined as the harmonic mean of these two values:

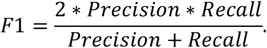

#### Cosine similarity

is a measure of similarity between two non-zero vectors defined in an inner product space. It is defined as the cosine of the angle between the two vectors, i.e., the dot product of the vectors divided by the product of their lengths. In the Type I2 simulation, the cosine similarity for each software is the cosine of the angle (denoted as *θ*) between the vector of true isoform proportion (denoted as 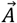) and the vector of estimated isoform proportion by the method (denoted as 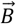). Similarly, in the Type F2. simulation, the cosine similarity for each method is the cosine of the angle between the vector of true fusion proportion and the vector of estimated fusion proportion by the method. The formula of cosine similarity is defined as

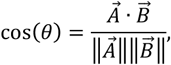

where *‖ · ‖* is *l*_2_-norm operator.

## Results

### IFDlong development

We developed software IFDlong, an isoform and fusion detector tailored for long-RNA-seq data. The software consists of six major steps, as shown in **Fig. 1C**.

### (Step 1) Long read alignment and filtering

IFDlong first takes in long-RNA-seq data and aligns long reads to a reference genome (e.g. GRCh38 for human and GRCm38 for mouse) using the noise tolerant long-read aligner Minimap2^29^. Unmapped long reads as well as reads with multiple alignments are filtered out.

### (Step 2) Gene, isoform and amino acid (AA) annotation

Long reads are intersected with gene/isoform annotation profile (described in Materials and Method for preparation) by BEDTools^30^. Reads with consecutive overlapping with known exons will be further annotated to known isoforms. While reads that cannot be fully covered by known exons will be defined as novel isoform. These reads will be further subject to gene-level annotation to be assigned into known genes or novel genes. Ultimately, all the long reads will be annotated three categories: known genes with known isoforms, known genes with novel isoforms, or novel genes. Note that, a long read will to be either uniquely assigned to one isoform or uncertainly aligned into multiple isoforms (**Fig. 1A**). As a special case, a long read is potentially aligned into more than one gene, which will serve as a supporting read for fusion transcript. Based on the transcript level annotation, IFDlong will predict the corresponding amino acid sequence using the AA annotation profile as reference (described in Materials and Method for preparation).

### (Step 3) Fusion transcript filtering

For the detection of fusion transcripts, false positives will be introduced due to nature of long-read sequencing (such as high error rate or artificial effect) and the complexity of the transcriptome (such as gene similarity and complex regulation mechanisms). To control the false positives, IFDlong applied the following filtering criteria. (1) Anchor length is defined as the number of base pairs that a supporting read is aligned to each fusion gene^23^. Short anchor length may result from misalignment, sequencing error or genome similarity. A long read is defined as a fusion supporting read by IFDlong only if it has a minimum of 10bp (by default) of anchor length to each fusion gene. (2) Fusion transcripts involving pseudo-genes are large likely to be false positives. (3) Fusion candidates with genes sourcing from the same family will be filtered out, because these are large likely result from multiple alignment due to the transcriptome similarity. Finally, a read-based report file will be generated to summarize the gene, isoform, amino acid, and fusion annotation. We would suggest the users to further filter the fusion transcripts will readthrough gene pairs or genes with short distance.

### (Step 4) Estimation of Isoform and fusion transcript expression

Based on the gene/isoform annotation report, IFDlong will estimate the isoform and fusion transcript expression. Per gene or fusion gene group, IFDlong estimates the relative abundance of isoforms and fusion transcripts by an Expectation-Maximization (EM) algorithm for those uncertain isoform assignments (for example, result from truncated long reads in **Fig 1A**, or fusion genes with multiple isoforms in **Fig 1B**). In the expectation step (E-step), IFDlong takes in read annotation information to calculate the likelihood of isoform/ fusion expression. And in the maximization step (M-step), the relative abundance will be calculated to optimize the likelihood function. The E-step and M-step will be performed iteratively until converge. The details were described in the Methods and Materials. Finally, IFDlong will output reports for isoform and fusion quantity estimates.

### Isoform annotation and quantification in simulation data

Multiple simulation data sets were generated to evaluate the pipeline performance in terms of isoform annotation and quantification. To mimic the real sequencing data, our IFDlong pipeline was first applied into the ONT long-RNA-seq data on Universal Human Reference (UHR) sample^12^ and quantified the expression intensities of 28157 isoforms (**Fig. S1A**). Based on this distribution of isoform expression, the Type I1 simulation datasets were generated by tool PBSIM2^31^ with different accuracy settings (accurate, PacBio and ONT), gradient number of long reads (around 0.5 million, 1 million, 2 millions, and 4 millions reads, see **Fig. S1B**), and gradient mean read length (200bps, 500bps, 800bps, 1000bps, 1200bps, 1500bps, 1800bps, and 2000bps, see **Fig. S2**). Simulation details were described in Materials and Methods.

As a representative of the Type I1 simulation, **Fig. 2A** shows the performance comparison in the simulation data with accurate sequencing, 0.5 million reads and 1500bps mean read length. The proposed IFDlong pipeline and three cutting-edge isoform quantification tools (LIQA^12^, TALON^14^, and Mandalorion^13^) were applied into the simulation data and compared with the true isoform distribution. The distribution histograms of isoform intensities are shown in the diagonal blocks of **Fig. 1A**, and all their pairwise distribution scatter plots and Spearman’s correlations are presented in the bottom and upper blocks, respectively. Among all the tools, the IFDlong achieves the highest correlation with the truth (0.83), while the other tools in general under-estimate the isoform expression (scatter plot under the diagonal when compared with the truth) or unable to capture a subset of the known isoforms (multiple zero-expressed isoforms in the scatter plot). The same conclusion holds for the simulation datasets with gradient parameter settings. **Fig. 2B and S3A** illustrate the Spearman’s correlation and root mean squared error (RMSE) in the three sequencing accuracy settings. The IFDlong reaches the highest performance followed by LIQA. In general, all the tools are robust to different platforms. The IFDlong shows slightly higher performance in the high-accurate setting compared with the ONT and PacBio settings. When checking the pipeline performance in terms of sequencing depth, **Fig. 2C and S3B** indicate that all the tools are robust to different sequencing coverage and are able to quantify isoform expression at relatively lower depth (such as 0.5M). The IFDlong presents slightly better performance along the increasing number of reads. In addition to sequencing accuracy and depth, tool performances were evaluated in gradient numbers of read length. As shown in **Fig. 1D and S3C**, all the tools yield dramatically increasing accuracy as the sequencing reads extending longer, which strongly prove the advantage of long-read techniques compared with short-read sequencing. Specifically, the IFDlong has already achieved high correlation (0.74) with the truth in the simulation data with 200bps mean length, and its performance further increases to 0.85 in 2000bps length.

**Figure 2:**
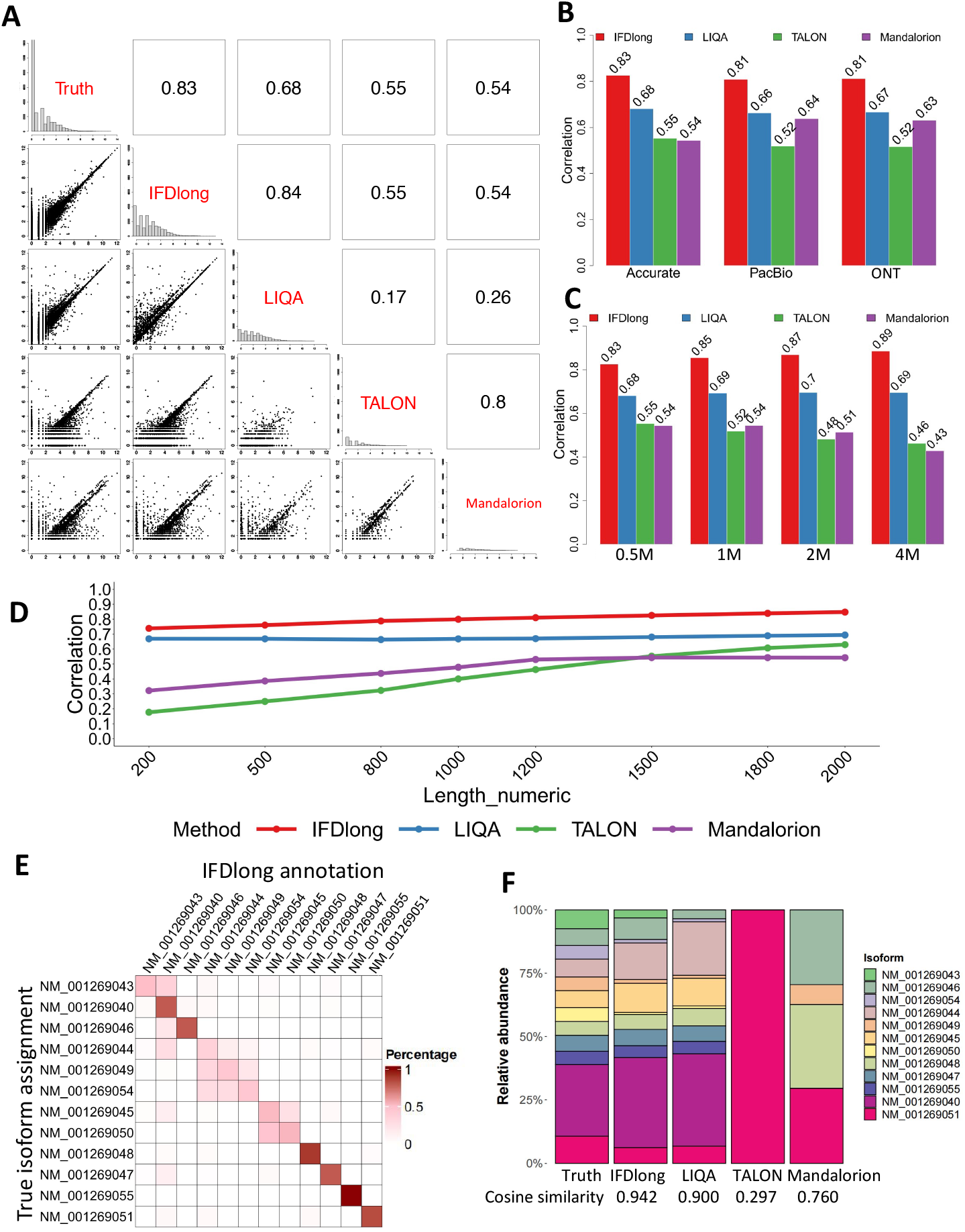
Isoform annotation and quantification in simulation analyses. (A) Pairwise comparison of the isoform quantification tools in the Type I1 simulation data with accurate-sequencing, 0.5M reads and 1500 bp median length. The bottom left cells present the pairwise scatter plot of isoform expression. The upper right cells indicate the Spearman’s correlation. (B) Spearman’s correlation of isoform expression between the truth and each tool under different sequencing accuracy in the Type I1 simulation with 0.5M reads and 1500 bp median length. (C) Spearman’s correlation of isoform expression between the truth and each tool under different sequencing depth in the Type I1 simulation with accurate sequencing and 1500 bp median length. (D) Spearman’s correlation of isoform expression between the truth and each tool under gradient median read lengths in the Type I1 simulation with 0.5M reads and accurate sequencing. (E)The percentage contingency table between the true isoform assignments and the annotation by IFDlong in the Type I2 simulation data. (F) Comparison of isoform relative abundance among the true simulations and the expression estimation by different tools in the Type-I2 simulation.

Besides the accuracy, computational costs were benchmarked among multiple tools in the meanwhile. In general, all the tools consume longer running time for larger sequencing coverage and longer read length with some exceptions (**Fig. S4A and S4B**). While the largest memory used is not a monotone function of the read length (**Fig. S4**), but all within acceptable cost. The IFDlong algorithm requires longer but manageable running time and similar memory cost compared with most of the other tools.

### Distinguishing multiple alternative splicing events sourcing from the same gene in simulation data

Other than gene-level quantification, one of the major advantages of long-read technique is to identify multiple alternative splicing events and quantify their expressions. As illustrated in **Fig. 1A**, a long read will be either uniquely aligned to one isoform or uncertainly assigned to multiple isoforms. To address this issue, the IFDlong pipeline developed the EM algorithm to estimate isoform relative abundance (see the Methods and Materials for algorithm details). In the Type I2 simulation, we generated long-reads on 16 isoforms sourcing from gene CTCFL as an example. Some isoforms were merged into one group given they share the same coding sequence (NM_001269040 includes NM_001269040, NM_001269041, NM_001269042, and NM_080618; NM_001269051 includes NM_001269051 and NM_001269052). Ultimately, in total 12 isoform groups were quantified for the performance evaluation. The isoform tools (IFDlong, LIQA, TALON and Mandalorion) were applied into this simulation data to quantify the abundance of all the alternative splicing events. Importantly, IFDlong and TALON are the only tools that can perform read-level isoform annotation, while the other pipelines only report the overall isoform abundancy/ intensity. **Fig. 2E** shows the contingency table (in percentage) between the true and the IFDlong annotated isoform assignments for the simulation data with 1500 bps median read length in high-accurate setting, where larger percentages in the diagonal block of the heatmap indicate higher consistency. In terms of abundance estimates, we compared the relative abundance (sum up to be 100%) among the truth and the estimates by the four tools in **Fig. 2F**. The IFDlong and LIQA show out the highest consistency with the truth (cosine similarity to be 0.94 and 0.90, respectively), followed by the Mandalorion and TALON. Similar as the Type I1 simulation setting, we simulated gradient read lengths in this Type I2 simulation setting. **Fig. S5** illustrates the contingency heatmap between the truth and the IFDlong pipeline with different read lengths. And the results show out increasing similarity when the read length extends longer.

### Novel isoform detection in simulation data

In addition to quantifying known isoforms, IFDlong can call novel isoforms from the long-RNA-seq data. As the toy example shown in **Fig. 3A**, four types of novel isoforms were defined: skipping exon, additional exon, alternative 5’ splice site, and alternative 3’ splice site. Multiple factors will cause the alterations of the reads compared with the reference sequences, such as insertion, deletion or mismatches caused by SNP, mutation and sequencing errors, or misalignment caused by multiple alignment and aligner performance. To avoid these false positives when calling novel isoforms, we proposed a buffer parameter for tolerance region next to the edge of the known exon. The buffer length is 9bps by default and can be adjusted by user setting. As shown in **Fig. 3B**, if the edge of the detected exon locates within the buffer region of the reference exon, the software will regard it as normal isoform, otherwise novel isoform with either truncation or extension. To simulate these novel isoform events, in the Type I3 datasets, we generated six libraries for each category of novel isoform. Each library contains half long-reads that are derived from normal isoforms, and another half reads supporting the novel isoforms of each category (details were described in Methods and Materials). Given only IFDlong and TALON can perform read-level isoform annotation, **Fig. 3C** compares the sensitivity and specificity between IFDlong and TALON in terms of detecting the long reads supporting the novel isoforms. Two buffer tolerance lengths were applied to IFDlong: 0bp (exact edge-matching between the reference and detected exon) and 9bps, while the buffer length for TALON is not allowed to be customized.

**Figure 3:**
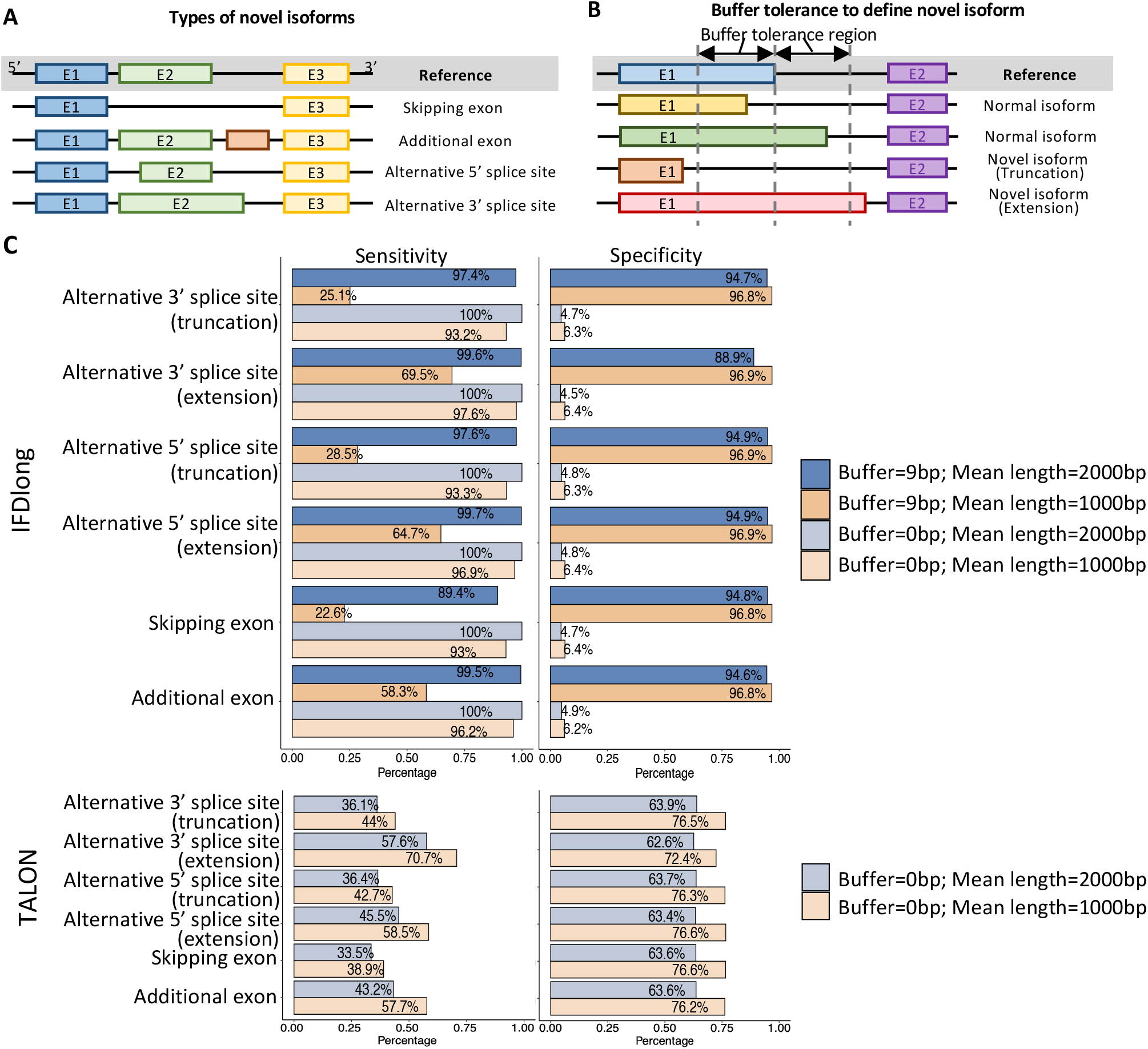
Illustration and detection of novel isoform. (A) Major types of novel isoforms. (B) Buffer tolerance to define novel isoform. Compared with the reference exon (blue), if the difference is within the buffer tolerance region, the isoform is still defined as normal one (yellow and green isoforms); otherwise, it will be defined as novel isoform with either truncation (orange) or extension (red). (C) Sensitivity (left) and Specificity (right) of IFDlong (top) and TALON (bottom) when detecting novel isoforms in the Type I3 simulation. The bars are colored by different buffer length setter (0bp or 9bp) and median length (1000 bp or 2000 bp) settings.

We are expecting that a reasonable buffer setting (suggested setting: 3-6bp) will dramatically control the false positives without missing a lot of true positives.

As shown in **Fig. 3C**, compared with zero buffer setting, IFDlong with 9 bps buffer region can significantly increase the specificity with reasonable sacrifice of sensitivity in high-accurate sequencing data. In this Type I3 simulation datasets, we generated datasets with mean read length to be 1000bps and 2000bps. As illustrated in **Fig. 3C**, IFDlong achieves higher performance in the longer read length datasets. For example, when detecting novel type of additional exon, IFDlong with 9bps buffer region can reach 99.5% sensitivity and 94.6% specificity, while the 1000bp can already reach satisfiable performance (e.g., with 9bps buffer, 58.3% sensitivity and 96.8% specificity for the novel type of additional exon). When compared with TALON, IFDlong with 9bps buffer region achieved higher sensitivity and specificity in all the 2000bps mean length datasets, and showed higher specificity and equivalent sensitivity in the 1000bps mean length datasets. In addition to high-accurate sequencing setting, simulation data with ONT and PacBio accuracy were generated, where similar conclusion holds in these two platforms (**Fig. S6**). Moreover, compared with the high-accurate sequencing reads in **Fig. 3C**, the 9bps buffer length benefits IFDlong more with significant increasing specificity and negligible sensitivity loss than 0bp buffer length when the sequencing error rate is high (**Fig. S6**). Overall, compared with TALON, our IFDlong pipeline is able to achieve higher sensitivity and specificity in most of the simulation scenarios (**Fig. 3C, S6** and **S7**).

### Application into real human datasets for isoform detection

Simulation data has its limitation to present the real sequencing. In order to evaluate the tool performance in a comprehensive manner, we applied the proposed IFDlong pipeline as well as the three existing isoform quantification tools into real sequencing data. For human data application, RNA extractions from both Universal Human Reference (UHR) and MCF7 human breast cancer cell line were sequenced by both ONT and PacBio platforms. Data were downloaded from public resources and fully described in Materials and Method. **Fig. 4A** and **S8** illustrates the Spearmen’s correlation of the true isoform distribution compared with the estimates by each one of the four tools. For UHR datasets, 964 isoforms quantified by TaqMan Real-Time PCR Assays is serving as the truth^32^. IFDlong achieved the highest correlation with the RT-PCR results (0.65 and 0.69 in ONT and PacBio dataset), indicating the high performance of our proposed tool. Among the 964 isoforms, the IFDlong has successfully captured 677 (70%) and 812 (84%) of them in ONT and PacBio dataset, respectively. Beyound these PCR quantified isoform set, in total 16578 and 28078 isoforms were detected by IFDlong pipeline in both datasets (**Table S1**), while the additional isoforms cannot be evaluated given the limitation of the RT-PCR measurement set. Similar analysis was performed on MCF7 datasets, where transcript expressions quantified by RNA-seq were used as underlying truth. As shown in **Fig. 4A** and **S8**, our IFDlong pipeline still performs the best compared with the other tools (Spearman’s correlation of 0.66 and 0.63 in both selected ONT and PacBio datasets). The full list of detected isoforms by IFDlong were summarized in **Table S2**.

**Figure 4:**
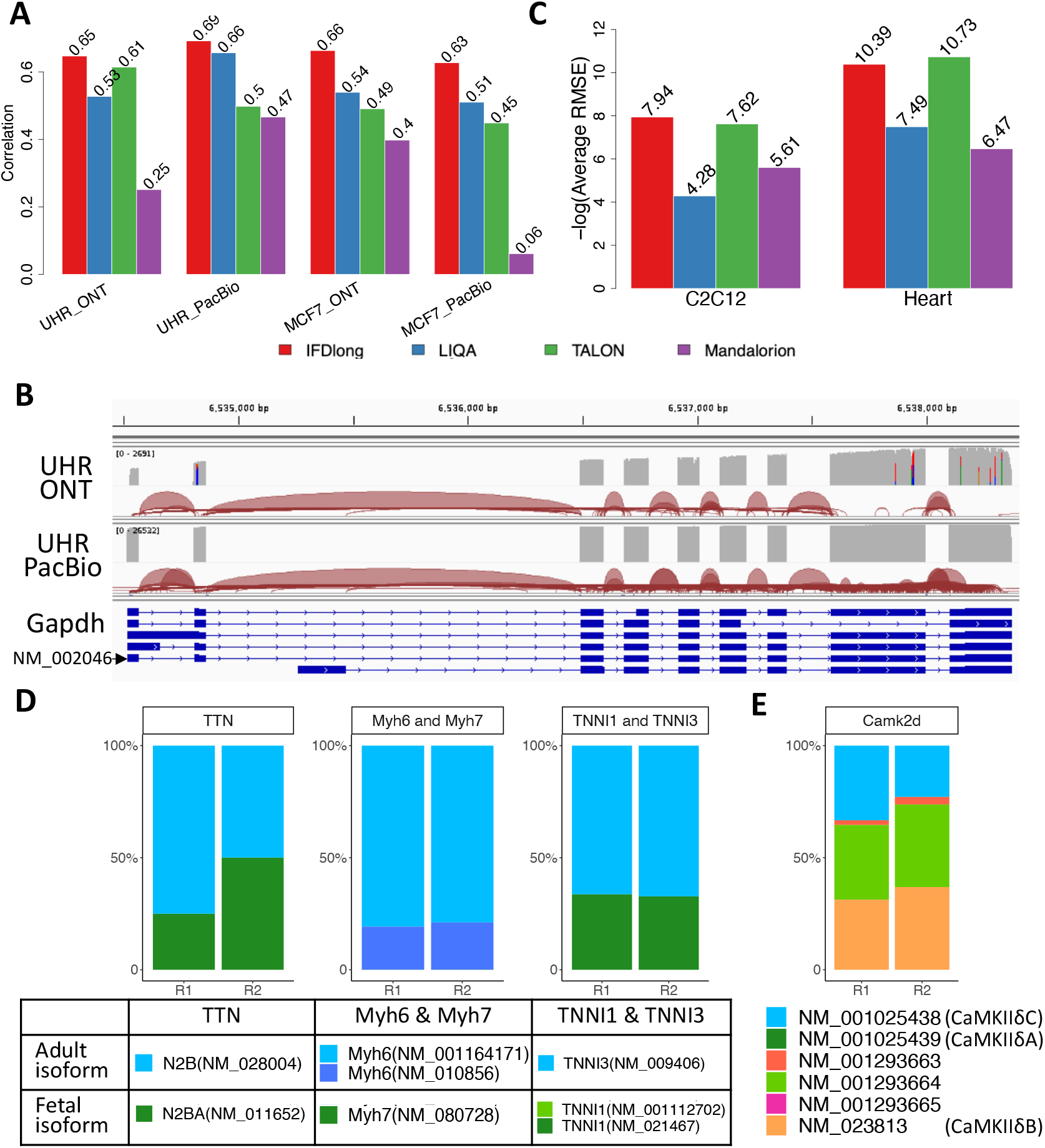
Application of isoform detection in real human and mouse studies. (A) Spearman’s correlation of isoform quantification in human studies: UHR sequenced by ONT and PacBio, and MCF7 breast cancer cell line sequenced by ONT and PacBio. (B) Illustration of multiple isoforms of gene Gapdh supported by long reads in UHR dataset. (C) Consistency measurement (-log10(average RMSE)) of isoform quantification in mouse studies: C2C12 cell line with four repeats and mouse heart tissues with two repeats. (D) Relative abundance of adult and fetal isoforms quantified by IFDlong pipeline on mouse heart tissues. (E) Relative abundance of multiple alternative splicing events of Camk2d quantified by IFDlong pipeline on mouse heart tissues.

As an illustration, **Fig. 4B** takes the house-keeping gene glyceraldehyde-3-phosphate dehydrogenase (Gapdh) as an example to show how the long-reads can quantify the transcript expression at isoform resolution. Gapdh is a highly expressed gene in human samples. Among the multiple alternative splicing events, NM_002046 plays a dominant role in the UHR datasets as quantified by IFDlong pipeline (**Table S1**). In both ONT and PacBio datasets, the long reads can successfully distinguish the differences in exon 1, 4, 6, and 7 among multiple isoforms.

### Application into mouse datasets for isoform detection

In addition to human data, our proposed pipeline can be applied into other species as long as the reference genome and annotation files are provided as the input. In this project, we’ve collected long-read data on mouse cell cline C2C12 (with four repeats) and heart tissues (with two repeats) from ENCODE project^33^ to illustrate the generalizability of our proposed tool into broad applications. For both datasets, TALON pipeline was employed by the ENCODE project to quantify the isoform expression. In addition, we applied IFDlong, LIQA and Mandalorion into these two mouse datasets for performance evaluation and application. Isoforms quantified by IFDlong were summarized in **Table S3 and S4**. For the mouse data, no golden standard truth is available, such as RT-PCR quantified isoforms or short-read RNA-seq measurements as a reference. Given this limitation, we can only check the performance similarity across the four tools. As shown in **Fig. S9**, IFDlong presents high agreement with LIQA and with TALON in all the datasets. In addition, the mouse datasets were applied to benchmark the consistency of isoform quantification in multiple repeats. Per software, mean isoform abundance was first calculated across multiple repeats, and then the average RMSE were derived based on this mean abundance. As shown in **Fig. 4C**, our IFDlong pipeline and TALON (result provided by ENCODE project^33^) achieves the largest -log10(average RMSE) value in both C2C12 and heart datasets, indicating the high robustness properties of the two tools.

Besides robustness, biological insights summarized from the isoform quantification can serve as indirect evaluation of the IFDlong pipeline. The heart tissues are collected from mouse at 14 days, which is the transition time of fetal isoforms to adult ones. The isoform quantification by our IFDlong pipeline (**Table S4**) has strongly supported this conclusion. For example, titin (TTN) is a massive sarcomeric protein serving as a molecular spring within the sarcomere, determining the passive stiffness of cardiomyocytes. During the perinatal period, there is a transition in titin isoforms from the compliant fetal titin N2BA (NM_011652) to the stiffer N2B (NM_028004) adult isoform in the heart, adapting to postnatal cardiac load demands^38^. Altered splicing of TTN is implicated in heart failure, and lowering the N2BA:N2B ratio has been proposed as a therapeutic strategy for heart failure^39-41^. The expression levels of N2BA and N2B that have been quantified by IFDlong presents this dynamic transition status (**Fig. 4D**). Besides TTN, the switching of several myofibril proteins during postnatal period are observed. For example, myosins are actin-based motor molecules with ATPase activity essential for muscle contraction. During perinatal period, the fetal isoform of myosin heavy chain 7 (Myh7, NM_080728) is gradually switching to adult isoform Myh6 (NM_001164171 and NM_010856)^42^. Similar isoform transition event was observed on troponin I (TNNI), which is the inhibitory subunit of troponin and the thin filament regulatory complex. During the postnatal period, the slow skeletal TNNI (TNNI1) transitions to the cardiac-type troponin (TNNI3)^43^. As shown in **Fig. 4D**, the IFDlong quantified isoforms have captured this transition process and more adult isoforms were observed than their corresponding fetal isoforms.

In addition, the mouse data also shows the distribution of multiple isoforms sourcing from the same gene. For instance, calcium/calmodulin dependent protein kinase II delta (CaMKIIδ) plays a central role in a variety of cardiac diseases^44^ [cite], which presents several alternative splicing forms as shown in **Fig. 4E**. CaMKIIδB (NM_023813) is a dominant isoform containing exon 14, which plays a critical role in gene transcription regulation and may play a protective role in cardiac diseases^45^. CaMKIIδC (NM_001025438) is another dominant isoform lacking exons 14-16, which is the major contributor of the pathological process of cardiac diseases^46^. CaMKIIδA (NM_001025439) comprises exon 13 and 15–17 and regulates L-type Ca2+ channel and contributes to the calcium mishandling in heart failure^47^. Besides CaMKIIδ, as shown in **Fig. S10**. the IFDlong analysis has revealed the isoform distribution of many other cardiac disease related genes, such as Calcium Voltage-Gated Channel Subunit Alpha1 C (Cacna1c), Troponin T2 (Tnnt2), Calcium/Calmodulin Dependent Protein Kinase II Gamma (Camk2g) and Myocyte Enhancer Factor 2C (Mef2c). All these have proved the additional resolutions that have been revealed by isoform-level analysis when compared with conventional gene-level quantification.

### Detecting differentially expressed isoforms in single-cell long-read RNA-seq data

The IFDlong pipeline is not only compatible with bulk long-RNA-seq, but also can be applied into single-cell long-RNA-seq data to investigate the isoform expression at single-cell resolution. A paired tumor-normal liver sample from a hepatocellular carcinoma (HCC) patient sequenced by LoopSeq was used in this project (details described in Materials and Methods)^34^. By IFDlong pipeline, in total 32,124 isoforms across 162 cells from the benign libraries and 285 cells from the tumor libraries were finally analyzed. Instead of the traditional gene-based analysis, we performed single-cell clustering and differential analysis based on the isoform profile. As shown in **Fig. 5A**, cells from tumor and normal libraries are well separable from each other. Due to the heterogeneity of the tumor and benign tissues, few cells sourced from the tumor library present benign expression patterns. Next, differential expression analysis was performed comparing tumor and normal cells to define differentially expressed isoforms (DEI) (**Table S5**). Fold changes of isoform were utilized for downstream Gene set enrichment analysis on KEGG database (**Fig. 5B** and **Table S6**) and GO database (**Fig. S11A** and **Table S6**). Top DEIs were subject to pathway enrichment test by Ingenuity Pathway Analysis (IPA on up-regulated DEI, **Fig. S11B** and **Table S6**). Top pathways primarily associated with the tumor immune microenvironment and immune evasion processes. These pathways play a crucial role in HCC tumorigenesis and in the response to immune checkpoint inhibitor (ICI) therapies. Particularly, the top pathways encompass various aspects of immune processes, including heterogeneity within the tumor microenvironment, immune cell infiltration, immune evasion mechanisms employed by tumor cells, and potential targets for immunotherapy (**Fig. 5B**).

**Figure 5:**
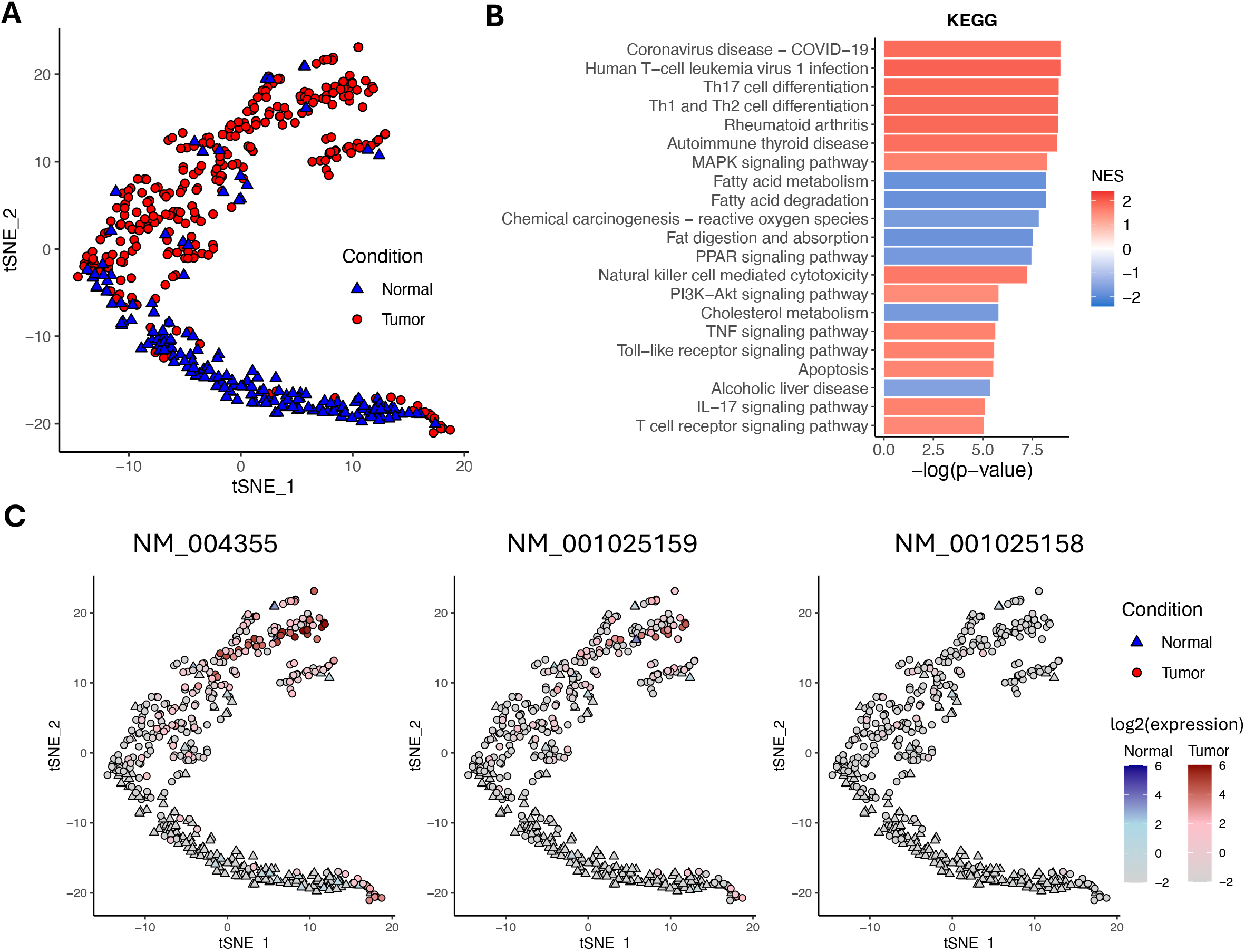
Application of IFDlong pipeline into single-cell long-read RNA-seq data on human HCC. (A)t-SNE visualization on the single-cell data based on isoform expression profile. (B) Top KEGG pathways enriched by the differentially expressed isoforms (DEI) comparing tumor and normal cells. (C) Multiple alternative splicing events of gene CD74 present differential expression patterns between normal and tumor cells.

For instance, considering the human leukocyte antigen (HLA)^48-50^ and B2M^51, 52^ genes, which encode proteins crucial for antigen presentation process critical for activating T cells and initiating an immune response against tumor cells. Dysregulation or alterations in the expression of these HLA genes can impact the immune system’s ability to recognize and eliminate cancer cells, directly influencing the success of ICIs^49, 51^. In addition, validated HCC immune players, CD74^53, 54^ (**Fig. 5C**), CCL5^55, 56^ (**Fig. S12A**), and IL7R^57, 58^ (**Fig. S12B**), known for their involvement in HCC ICI response, have been identified as top isoforms in pathway analysis. Their isoforms exhibit a significant increase in tumor cells compared to non-tumor regions, supporting the validated role of these isoforms in HCC pathogenesis, which underscores the necessity for understanding mechanisms of ICI resistance. Notably, CD74 contains three isoforms (**Fig. 5C**, NM_004355, NM_001025159, and NM_001025158). Based on IFDlong analysis, isoform NM_004355 and NM_001025159 present similar expression patterns, while NM_001025158 shows overall low expression. CD74 plays multifaceted roles in HCC pathogenesis by directly influencing antigen presentation through the regulation of major histocompatibility complex (MHC) class II molecules, influencing immune modulation and tumor cell behavior^53,54^. Furthermore, several isoforms of CCL5 (**Fig. S12A**, dominant by isoform NM_002985) and IL7R (**Fig. S12B**, with both NR_120485 and NM_002185 highly expressed) have been reported, resulting from alternative splicing or post-translational modifications^59-61^, may exert distinct biological functions in HCC, beyond their major roles as chemo-attractants and cognate receptors for various immune cell types, including T cells, monocytes, macrophages, and dendritic cells. Isoform diversity contributes to the establishment of an immunosuppressive microenvironment within the tumor, thereby facilitating tumor progression and immune evasion.

### Fusion isoform detection and quantification in simulation data

IFDlong is designed to detect and quantify fusion transcript at isoform resolution. For performance evaluation, the Type F1 dataset was generated by pooling reads simulated from 1000 normal isoforms and 1000 fusion transcripts. The proposed IFDlong pipeline and two existing fusion quantification tools (Genion^26^ and JAFFAL^27^) were applied into the simulation data and compared with the true fusion distribution. As a representative, **Fig. 6A** shows performance comparison in the data with approximately 0.5 million high-accurate sequencing reads. Considering that the Genion and JAFFAL can only detect fusion transcript at gene resolution, **Fig. 6A** illustrates the fusion gene expression in the diagonal blocks, and all their pairwise expression scatter plots and Spearman’s correlations in the bottom and upper blocks. Among all the tools, the IFDlong achieves the highest correlation with the truth (0.83), followed by JAFFAL (0.79) and Genion (0.75). The same conclusion holds for the simulation datasets with ONT and PacBio accuracy sequencing. As shown in **Fig. 6B and S13**, IFDlong always reaches the best performance for different accuracy settings. All the tools are generally robust to high sequencing errors, where the Spearman’s correlations for ONT and PacBio sequencing are equivalent to the ones for high-accurate sequencing.

**Figure 6:**
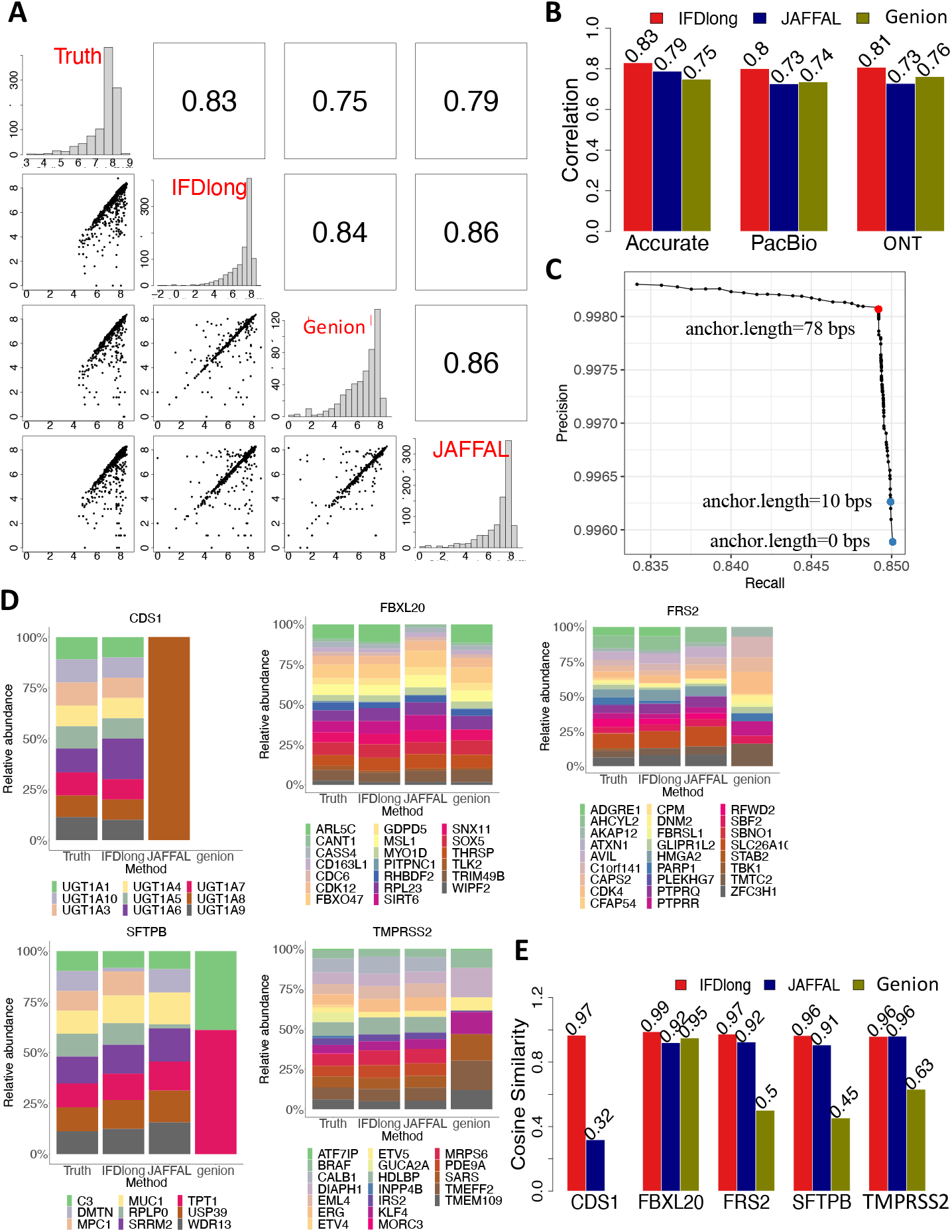
Detection and quantification of fusion transcripts in simulation studies. (A) Pairwise comparison of the fusion gene quantification tools in the Tpe-F1 simulation data with accurate sequencing. The bottom left cells present the pairwise scatter plot of fusion expression. The upper right cells indicate the Spearman’s correlation. (B) Spearman’s correlation of fusion expression between the truth and each tool under different sequencing accuracy in the Type F1 simulation data. (C) Cosine similarity of the relative abundance of multiple paired genes in the Type F2 simulation data. (D) Precision and Recall curve for fusion transcript detection by IFDlong pipeline using different anchor lengths. (E) Relative abundance of multiple paired genes in the Type F2 simulation data.

Exploring fusion transcript at isoform resolution is one of the major advantages of IFDlong. Compared with Genion and JAFFAL, IFDLong is the only tool that can annotate and quantify fusion isoforms (**Fig. 1B**). The scatter plots comparing the true fusion isoform intensities and the IFDlong estimated expression were shown in **Fig. S14**, where IFDlong can reach Spearman’s correlation of 0.47, 0.48 and 0.50 in the high-accurate, PacBio and ONT dataset, respectively. Given the increasing complexities of fusion quantification at isoform resolution compared with gene-level analysis, some fusion isoforms were under-estimate by IFDlong (dots locate underneath the diagonal in **Fig. S14**). One of the key factors for fusion detection is the setting of anchor length, which indicates the minimum base pairs of read alignment to each fusion gene^23^. Shorter anchor length resulted in high recall rate, but with the sacrifice of precision, because larger number of false positives with short aligned supporting reads were introduced. In contrast, longer anchor length can reach highest precision with the loss of recall given true fusions with shorter alignment will be mis-detected (large false negative). To test the tool sensitivity to anchor length, **Fig. 6C** illustrates the precision and recall (PR) curve of fusion transcript detection with anchor length ranging from 0bp to 100bps. Within this range, IFDlong can reach both high precision and recall rates (precision > 0.99 and recall > 0.83). **Fig. 6D** highlights the highest F1-score at anchor length to be 78bps. In addition to this high-accurate sequencing data (**Fig. 6D**), RR curves were drawn in the Type F1 simulation data with PacBio and ONT sequencing accuracy (**Fig. S12**). Decreasing sequencing accuracy resulted in comparatively lower precision recall rates, and required higher anchor length to reach the best F1-score. In practice, to discover more fusion candidates, a relatively small anchor length is suggested. IFDlong sets 10bps by default, but users can adjust this parameter to better balance the precision and recall and adapt to the real data.

### Distinguishing multiple fusion-mates paired with the same gene in simulation data

A popular gene may potentially pair with multiple mates in real practice. The Type F2 datasets were simulated to estimate the software performance in distinguishing different fusion-mates. Specifically, based on the TCGA fusion database, top five popular genes (FBXL20, CDS1, TMPRSS2, FRS2 and SFTPB) with their corresponding mate gene panels (**Table S7**) were selected as template to generate the simulation data. IFDlong, Genion and JAFFAL were then applied into this dataset to quantify the expression level of all fusion pairs. The relative abundances of all the fusion mates were compared across the truth and the software estimations in **Fig. 6D**, and their corresponding cosine similarities were summarized in **Fig. 6E**. For example, TMPRSS2 is a popular fusion gene in prostate cancer. Other than the most prevalent TMPRSS2-ERG fusion^62^, in total 19 genes were reported to be fused with TMPRSS2 in TCGA database (**Table S7**). Compared with the truth, both IFDlong and JAFFAL achieved high agreement (cosine similarity = 0.96) when estimating the relevant abundance of these 19 fusion transcripts in the simulation data with high-accurate sequencing and mean read length to be 1500bp (**Fig. 6D and 6E**). In all the five simulations, IFDlong presented the highest cosine similarity with the truth, while JAFFAL and Genion resulted in unrobust performance. All the three tools showed high consistency with the truth in FBXL20 simulation, but JAFFAL and Genion failed in CDS1 simulation. Similar results were observed in the simulation data with different accuracy settings (accurate, PacBio and ONT) and mean read lengths (200bp, 1000bp and 1500bp) (**Fig. S16-S18**). IFDlong achieved the highest cosine similarity (higher than 0.94) for all the simulation scenarios, followed by JAFFAL which performed well in all the settings other than CDS1.

### Two-segment and three-segment fusion detection in real data application

Besides *in silico* simulation datasets, the fusion callers were applied into real sequencing data for performance evaluation and novel fusion transcripts detection. The IFDlong fusion detection pipeline, as well as two existing tools (Genion and JAFFAL) were evaluated in four real data sets: MCF7 breast cancer (bulk long-RNA-seq by PacBio and ONT), H838 Lung adenocarcinoma cell line (single-cell long-RNA-seq by ONT), samples from colon cancer patients (bulk long-RNA-seq by LoopSeq), and liver tissue samples from hepatocellular carcinoma (HCC) patients (single-cell long-RNA-seq by LoopSeq). Among them, colon cancer and HCC cohorts are in-house samples, where novel fusions were detected and validated in our previous research^6, 34^ As shown in **Table 1**, The IFDlong pipeline successfully detected all these two-segment fusions, while Genion and JAFFAL can only identify 10/11 and 5/11 fusions. All the fusion callers were applied into MCF7 and H383 datasets and the two-segment fusions identified by IFDlong were listed in **Table S8**.

**Table 1:**
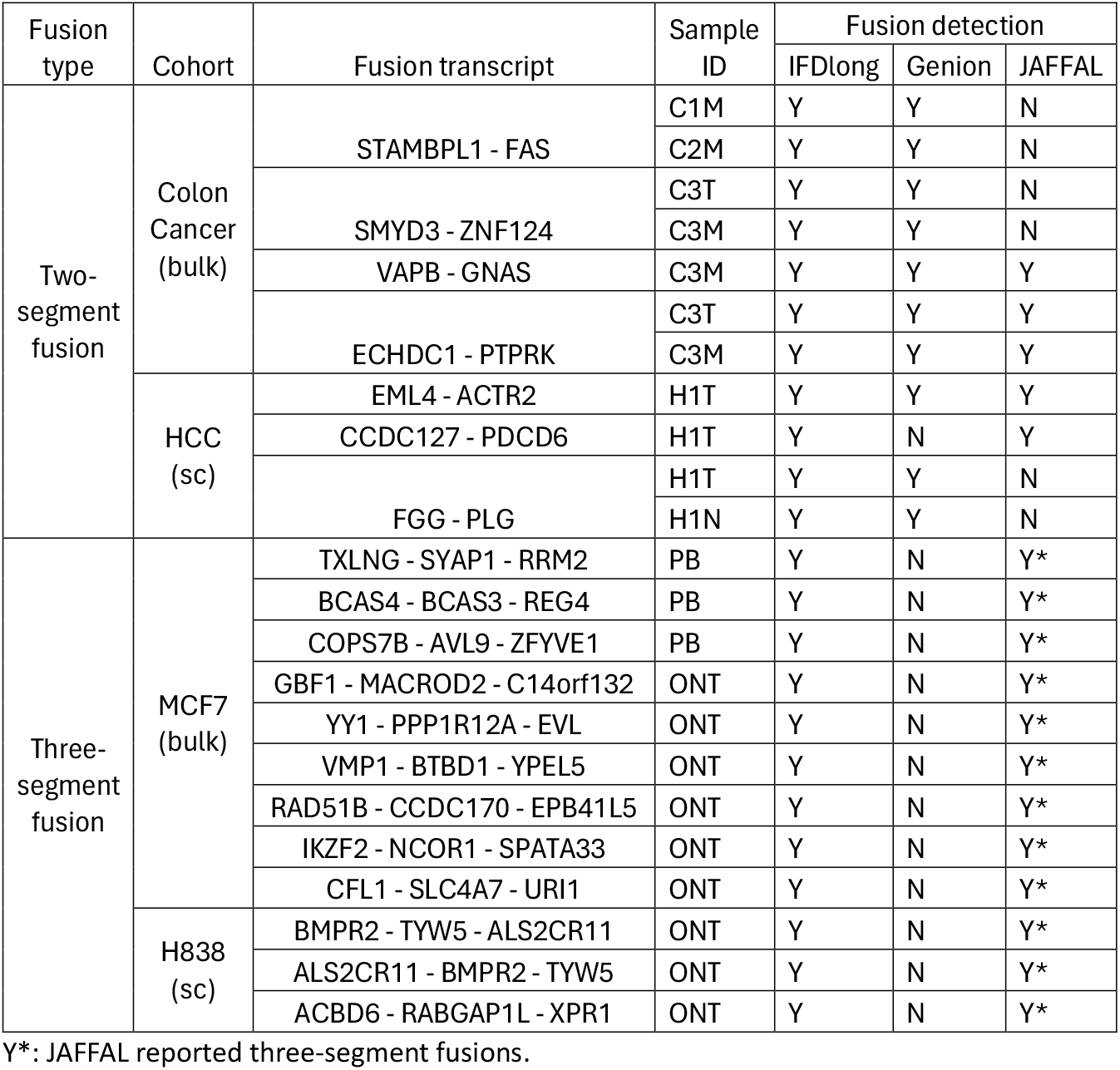
Two-way and three-way fusion detection in real data application.

| Fusion type | Cohort | Fusion transcript | Sample ID | Fusion detection |  |  |
| --- | --- | --- | --- | --- | --- | --- |
|  |  |  |  | IFDlong | Genion | JAFFAL |
| Two-segment fusion | Colon Cancer (bulk) | STAMBPL1 - FAS | C1M | Y | Y | N |
|  |  |  | C2M | Y | Y | N |
|  |  | SMYD3 - ZNF124 | C3T | Y | Y | N |
|  |  |  | C3M | Y | Y | N |
|  |  | VAPB - GNAS | C3M | Y | Y | Y |
|  |  | ECHDC1 - PTPRK | C3T | Y | Y | Y |
|  |  |  | C3M | Y | Y | Y |
|  | HCC (sc) | EML4 - ACTR2 | H1T | Y | Y | Y |
|  |  | CCDC127 - PDCD6 | H1T | Y | N | Y |
|  |  | FGG - PLG | H1T | Y | Y | N |
|  |  |  | H1N | Y | Y | N |
| Three-segment fusion | MCF7 (bulk) | TXLNG - SYAP1 - RRM2 | PB | Y | N | Y* |
|  |  | BCAS4 - BCAS3 - REG4 | PB | Y | N | Y* |
|  |  | COPS7B - AVL9 - ZFYVE1 | PB | Y | N | Y* |
|  |  | GBF1 - MACROD2 - C14orf132 | ONT | Y | N | Y* |
|  |  | YY1 - PPP1R12A - EVL | ONT | Y | N | Y* |
|  |  | VMP1 - BTBD1 - YPEL5 | ONT | Y | N | Y* |
|  |  | RAD51B - CCDC170 - EPB41L5 | ONT | Y | N | Y* |
|  |  | IKZF2 - NCOR1 - SPATA33 | ONT | Y | N | Y* |
|  |  | CFL1 - SLC4A7 - URI1 | ONT | Y | N | Y* |
|  | H838 (sc) | BMPR2 - TYW5 - ALS2CR11 | ONT | Y | N | Y* |
|  |  | ALS2CR11 - BMPR2 - TYW5 | ONT | Y | N | Y* |
|  |  | ACBD6 - RABGAP1L - XPR1 | ONT | Y | N | Y* |

Y\*: JAFFAL reported three-segment fusions.

In addition to two-segment fusions that are frequently observed, three-segment fusion were discovered in the previous research^63^, while majority of the predicted three-segment fusions may result from artificial chimera events or mis-alignment. To test the ability of three-segment fusion discovery, **Table 1** lists 12 fusion transcripts detected from MCF7 and H838 by JAFFAL^27^. The IFDlong pipeline can successfully detect all of them, while Genion failed to report three-segment fusions.

## Discussion

In this paper, we introduced IFDlong, a novel bioinformatics tool tailored for the analysis of long-RNA-seq data. IFDlong offers several distinct advantages over existing tools, making it a comprehensive solution for accurate annotation and quantification of isoforms and fusions. First, IFDlong offers a suite of functions that encompass various aspects of long-RNA-seq analysis (**Fig. 1C**), including long-read annotation at both gene and isoform resolution (**Fig. 2**), prediction of amino acid sequences, known isoform quantification (**Fig. 2**), novel isoform discovery (**Fig. 3**), and detection of fusion transcripts (**Fig. 6**).

Second, IFDlong performs statistical estimation to accurately infer the isoform and fusion abundance. Specifically, the EM algorithm was developed to address the long reads with ambiguous annotations (reads with uncertain isoform assignments, **Fig. 1A**) and fusion transcripts consist of multiple alternative splicing variants (**Fig. 1B**). IFDlong shows higher accuracy in terms of isoform and fusion quantification when compared with the existing tools (**Fig. 2 and 6**).

Third, IFDlong employs multiple selection criteria to control false positives in the detection of novel isoforms and fusion transcripts. IFDlong enhances the accuracy of fusion detection by filtering out fusion candidates involving pseudogenes, genes from the same family, and readthrough events. Moreover, the user-adjustable parameters, such as buffer length for novel isoform detection (**Fig. 3**) and anchor length for fusion identification (**Fig. 6**), provide flexibilities in customizing the analysis pipeline to adapt to specific experimental conditions.

Fourth, IFDlong presents advantages in its versatility and compatibility with diverse experimental setups and species. Specifically, IFDlong is applicable for both human and mouse data (**Fig. 4**), and can be easily generalized to other species given the corresponding reference and annotation profiles. Moreover, IFDlong is compatible with different experimental platforms (PacBio, ONT and linked-short-read platforms) and multiple library preparation strategies (bulk and single-cell RNA-seq, as shown in **Fig. 5**; direct RNA and cDNA libraries, as shown in Fig. S8). In addition, IFDlong can take in the alignment file by different long-read aligners, such as minimap2^29^ by default or STAR-long^64^. All these properties make IFDlong a generalizable tool that can be applied for broad applications and adjusted to best suits individual dataset.

In this project, comprehensive comparisons against existing tools were performed to demonstrate the superior performance of IFDlong through extensive simulations and real-data applications. Five types of simulation datasets were generated to prove the high performance of IFDlong in terms of isoform quantification (Type I1 and I2), novel isoform detection (Type I3), and fusion quantification (Type F1 and F2). Multiple real datasets were employed to test the generalizability of the IFDlong pipeline. The robustness and reliability of IFDlong make it a prioritized tool for researchers seeking accurate and comprehensive analysis of long-RNA-seq data.

Our analysis pipeline also highlights the significant potential of the single-cell long-RNA-seq data analysis. The HCC study enables the delineation of multiple isoform components at the single-cell resolution. This approach facilitates the identification of rare cell populations, evaluation of cell-to-cell variability, and the reconstruction of cellular trajectories, thereby offering a comprehensive view of the immune landscape in HCC. These approaches can be generalizable to comprehend the molecular mechanism underlying diverse disease and biological processes. These insights guide the development of more targeted and effective therapeutic strategies tailored to individual patients.

Admittedly, this study contains limitations to be addressed. In the comparison with other existing tools, the simulation data and the IFDlong pipeline were using the same hg38 annotation profile that was downloaded from UCSC Genome Browser. If the existing tools can be built from the user-defined annotation file, we will apply the same profile. If not, the embedded annotation file may not be the same as the one that is used by IFDlong, which will potentially cause mismatching due the different number of genes or alternative gene names in different databases. While for the comparison using real datasets, the qPCR or short-read RNA-seq data served as the truth, avoiding the annotation bias issue. For the performance comparisons, all the tools applied their default parameter settings, including our IFDlong pipeline. We admit that tuning parameters may influence the software performance, but this is out of the scope of this project.

Given all the advantages of the IFDlong, further improvements can be made as the future work. First, we will further increase the computing efficiency of the proposed software to make the running time and memory cost to be manageable for larger dataset. In addition, the current pipeline is based on the alignment profile by a single aligner (Minimap2 by default). To take the advantages of multiple aligners (such as STAR-long^64^, GMAP^65^ or other long-read aligners), the pipeline can be further improved by integrating multiple alignment files by more than one aligners.

In conclusion, IFDlong presents significant advancements in long-RNA-seq analysis for the annotation and quantification of isoforms and fusion transcripts. Its unique features, including the integration of the EM algorithm, stringent false-positive control, compatibility, and superior performance, position IFDlong as a versatile tool for long-read transcriptome research. This novel bioinformatics software will contribute to the community by its broad application into biomedical research.

## Funding

This research is supported in part by the Pittsburgh Liver Research Centre through NIH/NIDDK Digestive Disease Research Core Center grant P30DK120531 (Pilot and Feasibility Grants to S.L., Genomics and Systems Biology Core), the National Institutes of Health through Grant Number UL1TR001857 (Pilot Grant to S.L.), the Innovation in Cancer Informatics Discovery Grant (to S.L. and S.K.), the NIH funding R01CA258449 (to S.K.), and the University of Pittsburgh Center for Research Computing through the NIH award S10OD028483.

## Supplementary Figure Legend

**Figure S1:**
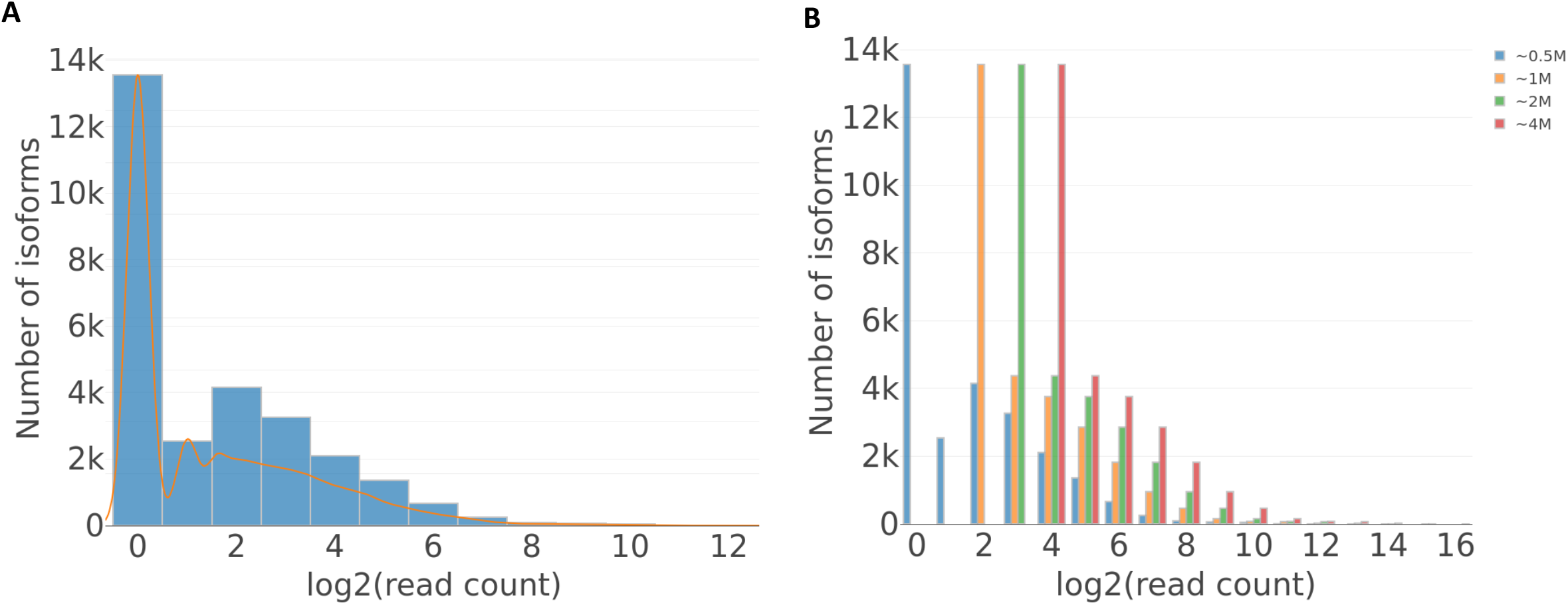
Read count distribution for the (A) Universal Human Reference real data and the (B) The Type I1 simulation data with gradient depth.

**Figure S2:**
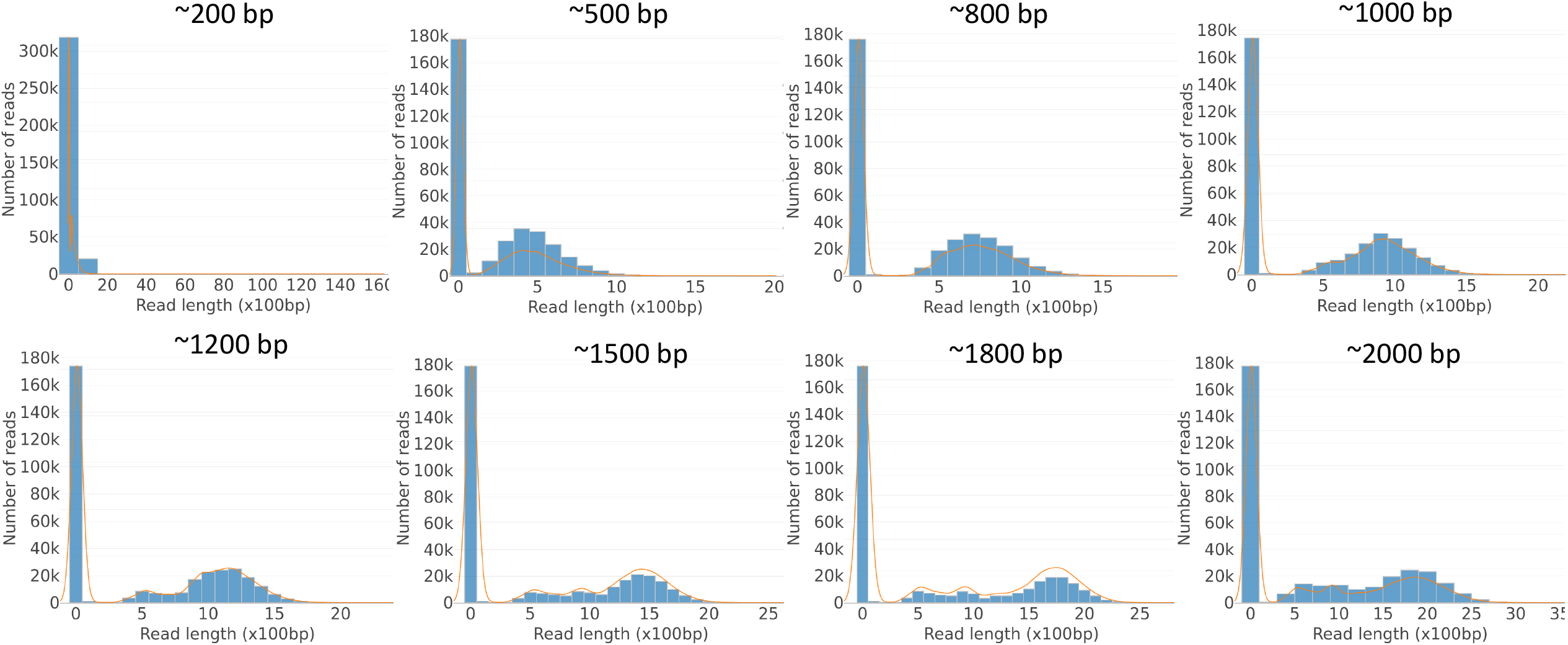
Read length distribution of the Type I1 simulation data with gradient median read length.

**Figure S3:**
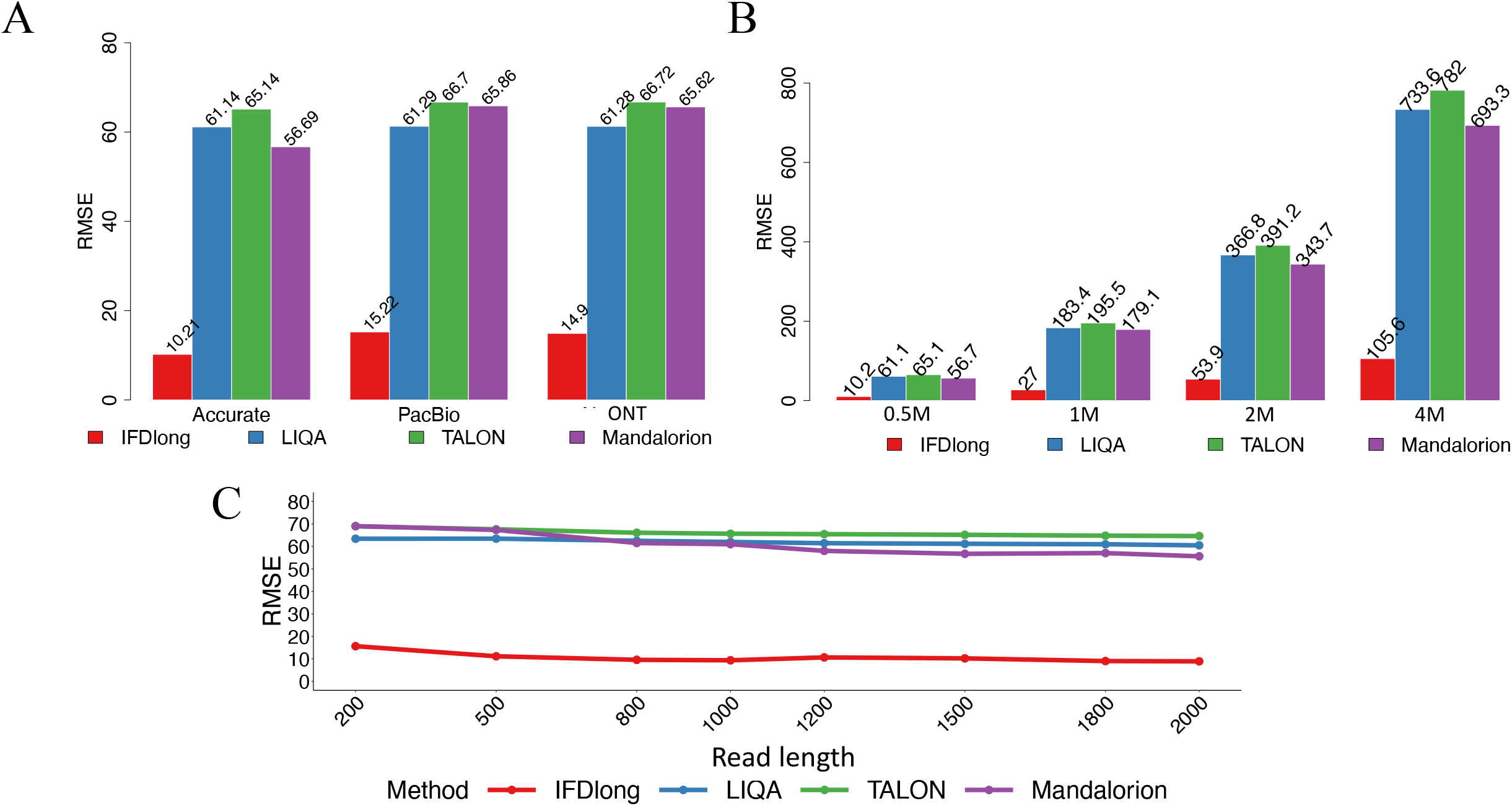
Root mean squared error (RMSE) between the true and the estimated isoform expression in the Type I1 simulation data. (A) RMSE of isoform expression between the truth and each tool under different sequencing accuracy with 0.5M reads and 1500 bp median length. (B) RMSE of isoform expression between the truth and each tool under different sequencing depth with accurate sequencing and 1500 bp median length. (c) RMSE of isoform expression between the truth and each tool under gradient median read lengths with 0.5M reads and accurate sequencing.

**Figure S4:**
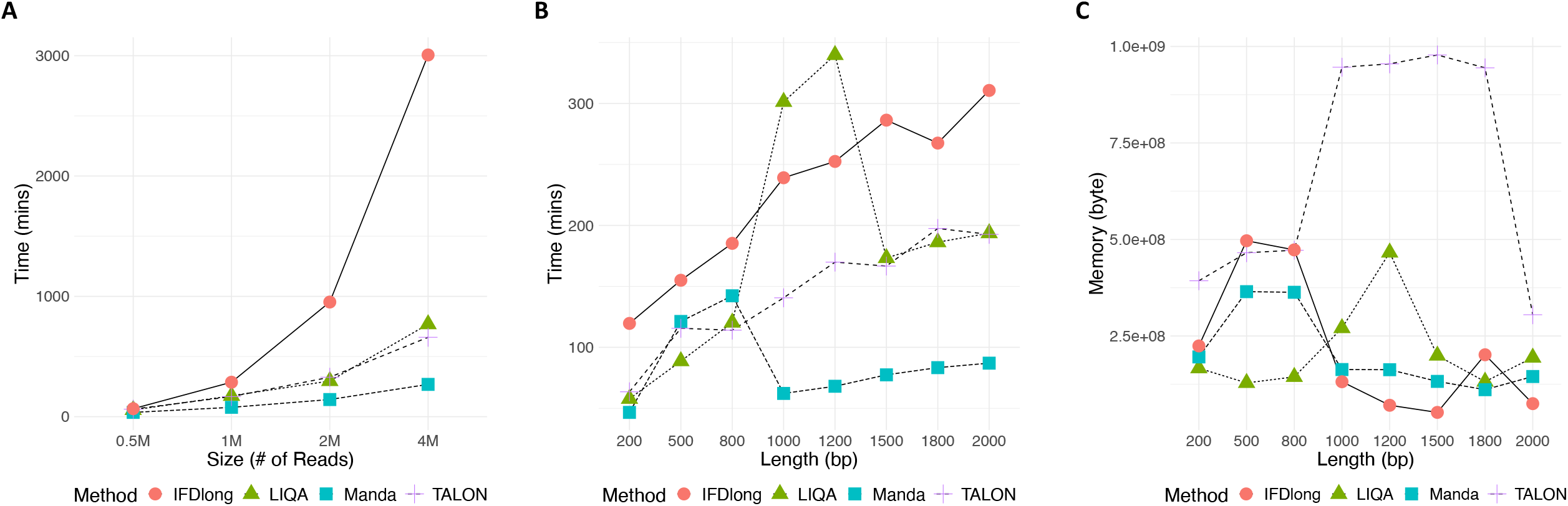
Computational cost for isoform quantification tools in the Type I1 simulation. (A&B) Time cost along different number of reads and read lengths. (C) Memory cost across different read lengths when working on 4M reads.

**Figure S5:**
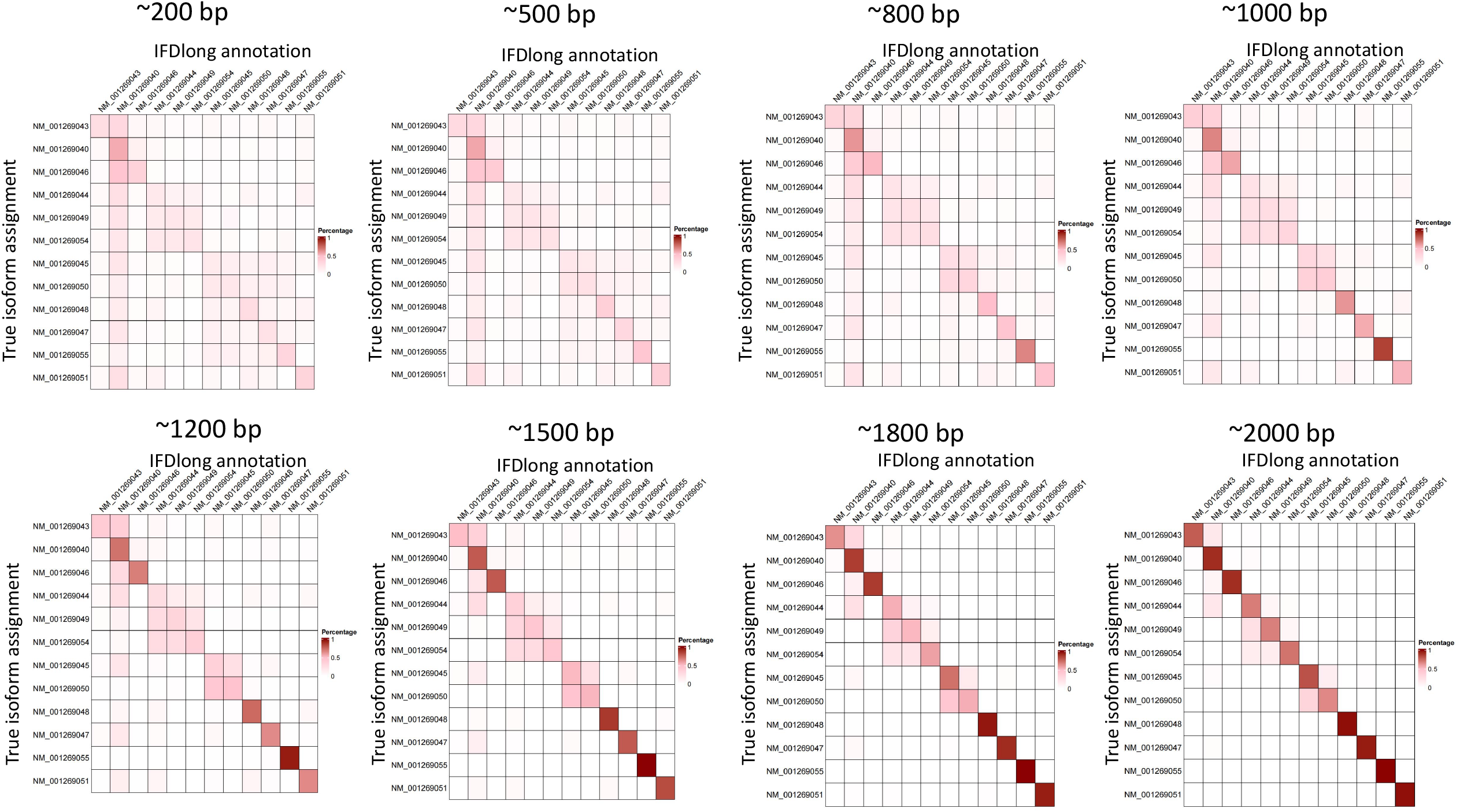
The percentage contingency table between the true isoform assignments and the annotation by IFDlong in the Type I2 simulation data with gradient median read lengths.

**Figure S6:**
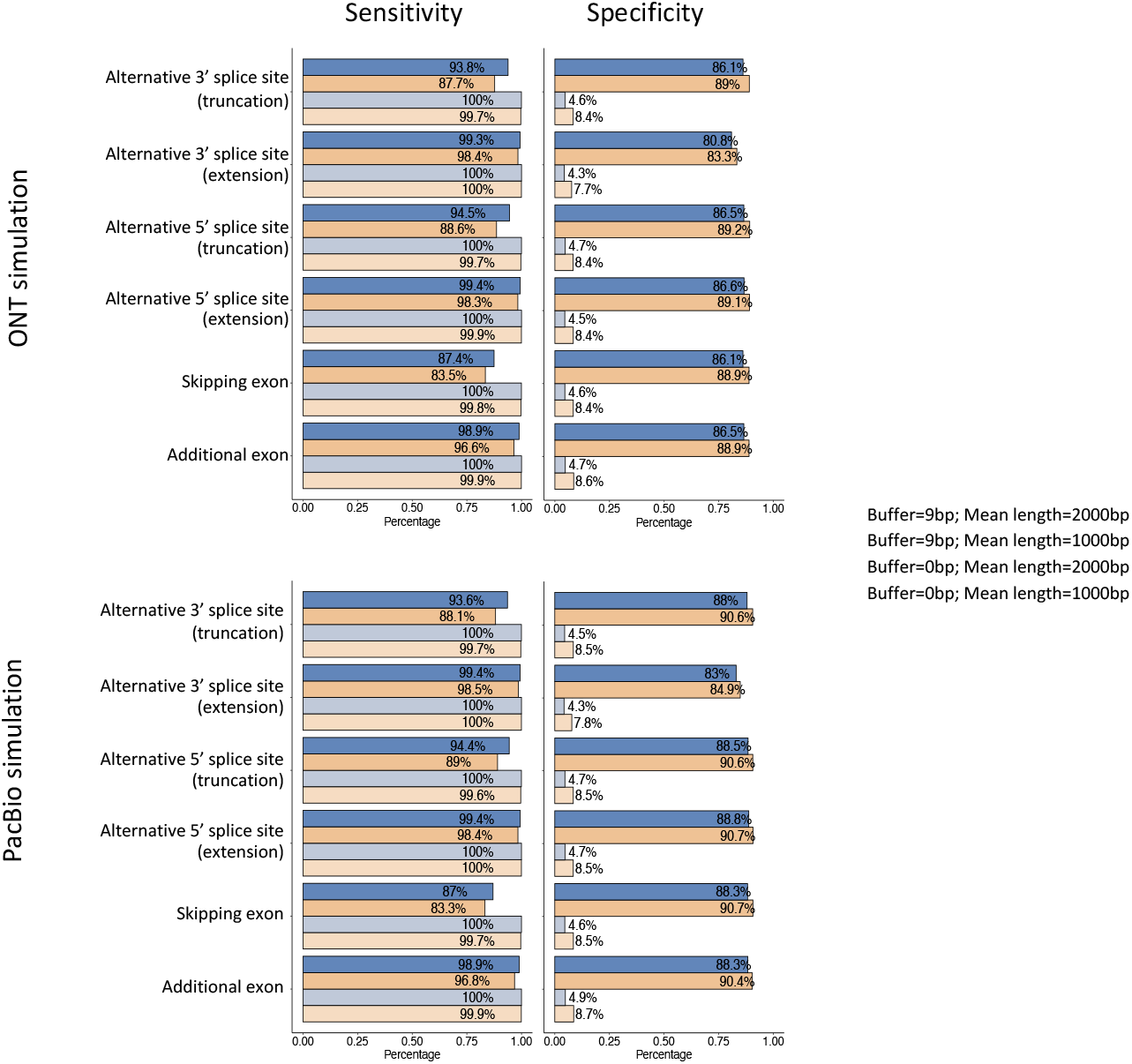
Sensitivity (left) and Specificity (right) of IFDlong when detecting novel isoforms in the Type I3 simulation with ONT (top) and PacBio (bottom) accuracy. The bars are colored by different buffer length setter (0bp or 9bp) and median length (1000 bp or 2000 bp) settings.

**Figure S7:**
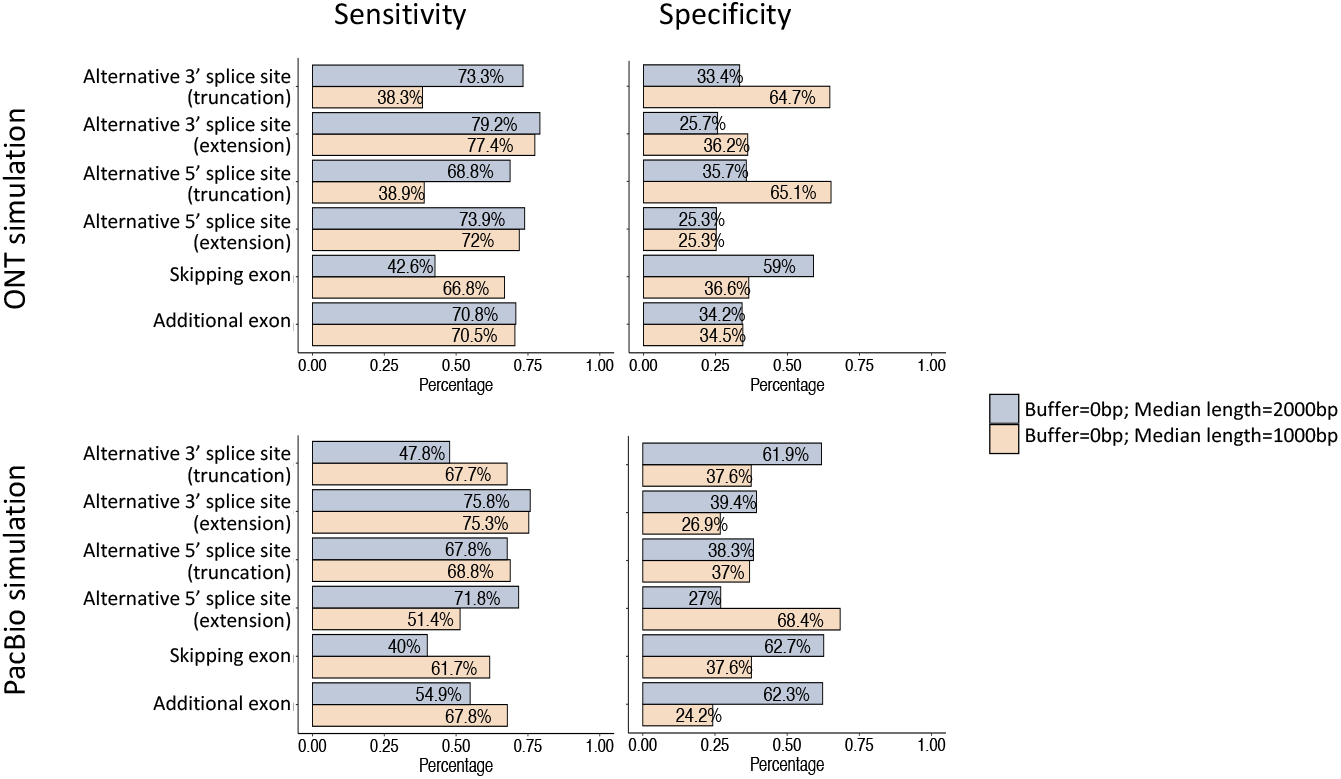
Sensitivity (left) and Specificity (right) of TALON when detecting novel isoforms in the Type I3 simulation with ONT (top) and PacBio (bottom) accuracy. The bars are colored by different median length (1000 bp or 2000 bp) settings.

**Figure S8:**
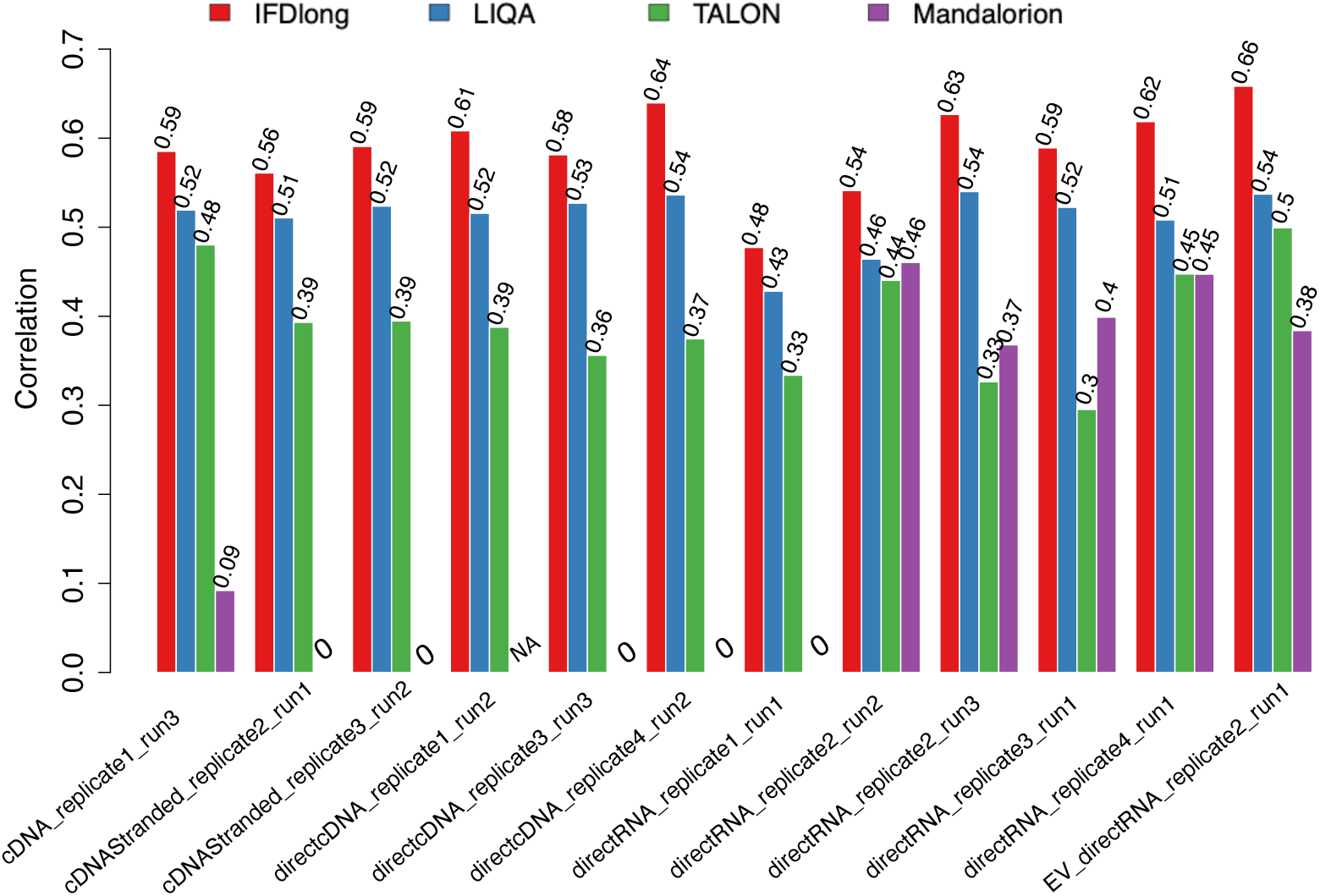
Spearman’s correlation of isoform quantification in human MCF7 breast cancer cell line by different ONT sequencing settings.

**Figure S9:**
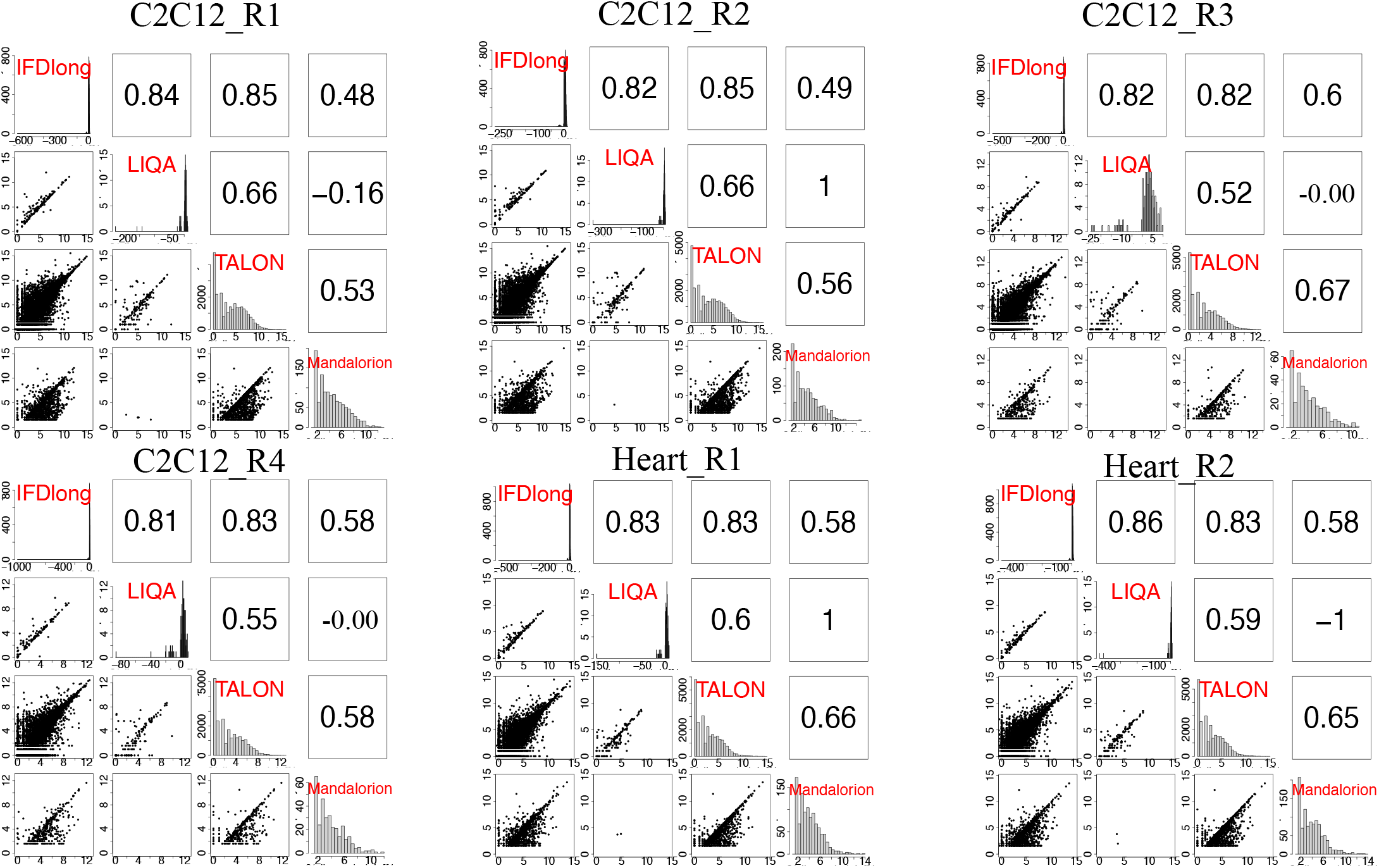
Pairwise comparison of isoform quantification in mouse C2C12 and Heart samples. The bottom left cells present the pairwise scatter plot of isoform expression. The upper right cells indicate the Spearman’s correlation.

**Figure S10:**
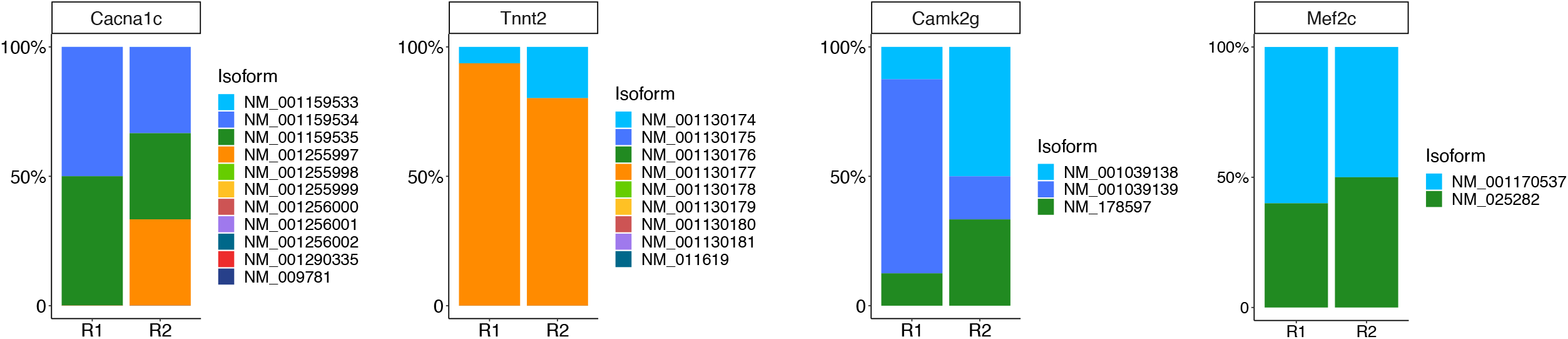
Isoform distribution of Calcium Voltage-Gated Channel Subunit Alpha1 C (Cacna1c), Troponin T2 (Tnnt2), Calcium/Calmodulin Dependent Protein Kinase II Gamma (Camk2g) and Myocyte Enhancer Factor 2C (Mef2c) in heart tissue.

**Figure S11:**
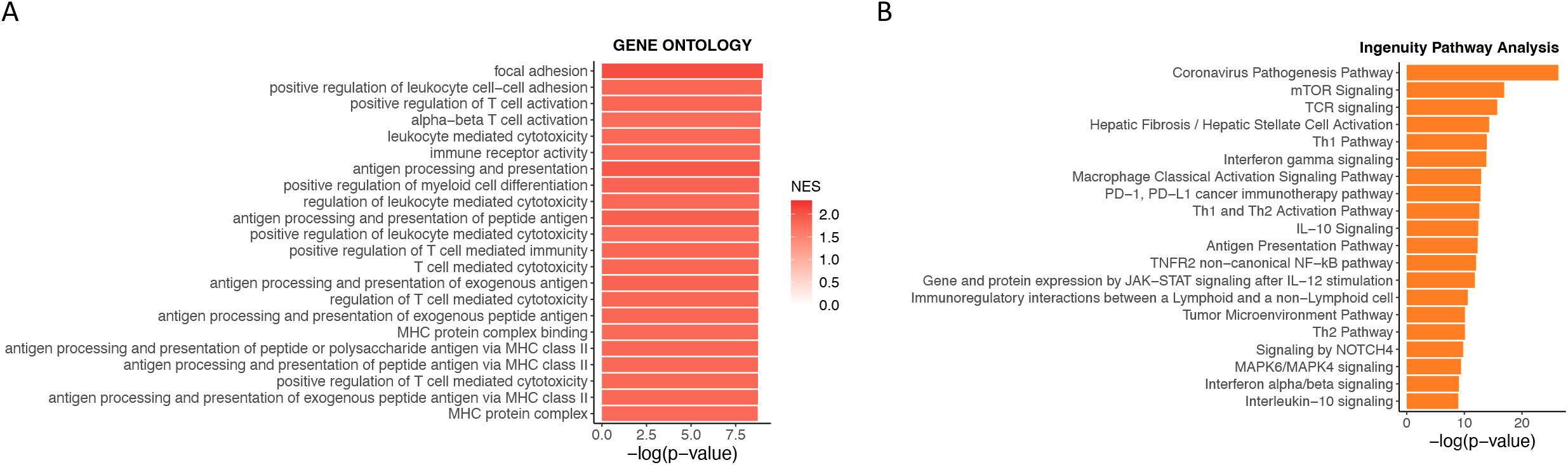
Pathway analysis on isoform differential expression analysis in HCC data. (A) Gene set enrichment analysis (GSEA) on Gene ontology (GO) database. (B) Ingenuity pathway analysis (IPA).

**Figure S12:**
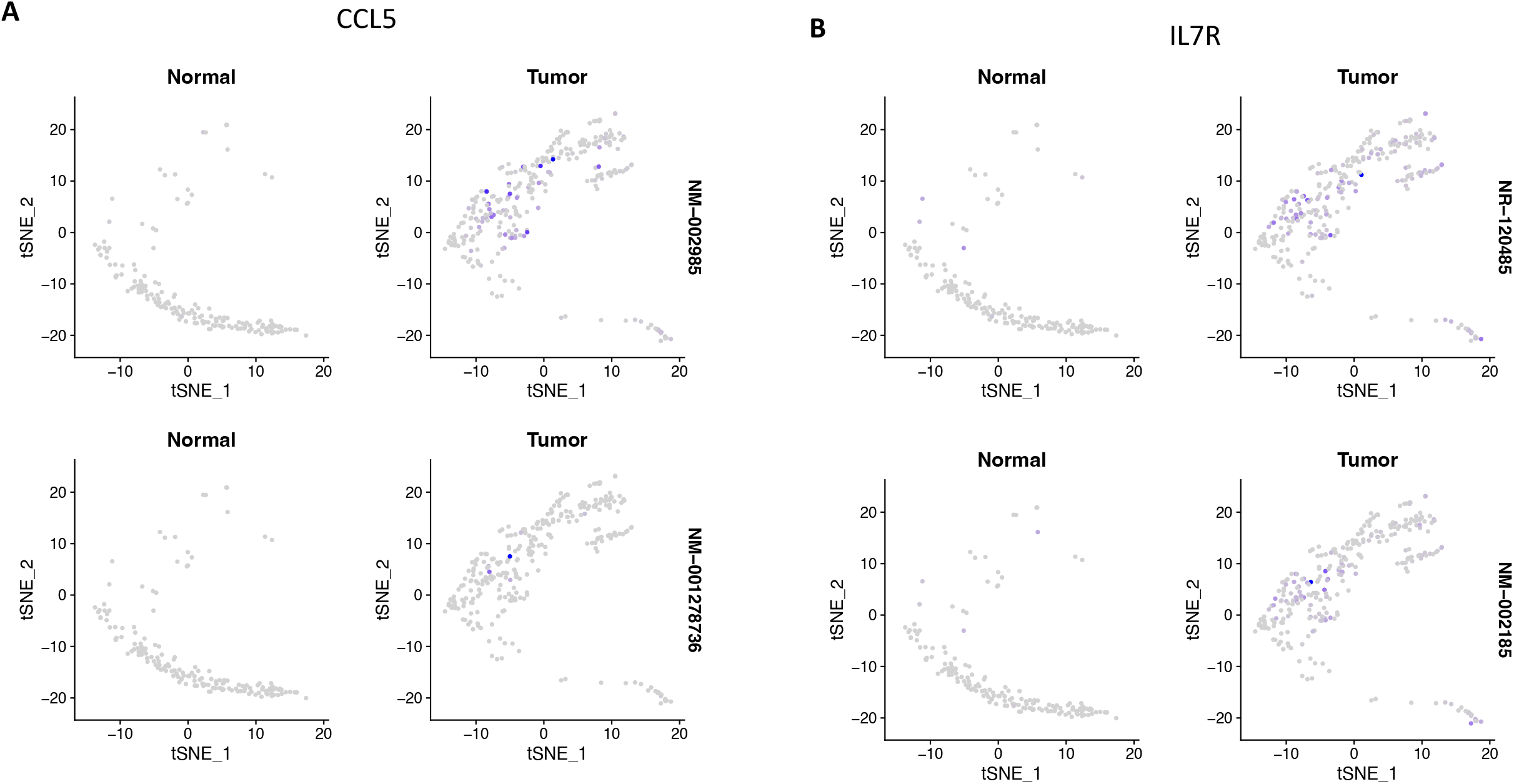
Feature plot of isoform expression for gene CCL5 and IL7R in HCC data.

**Figure S13:**
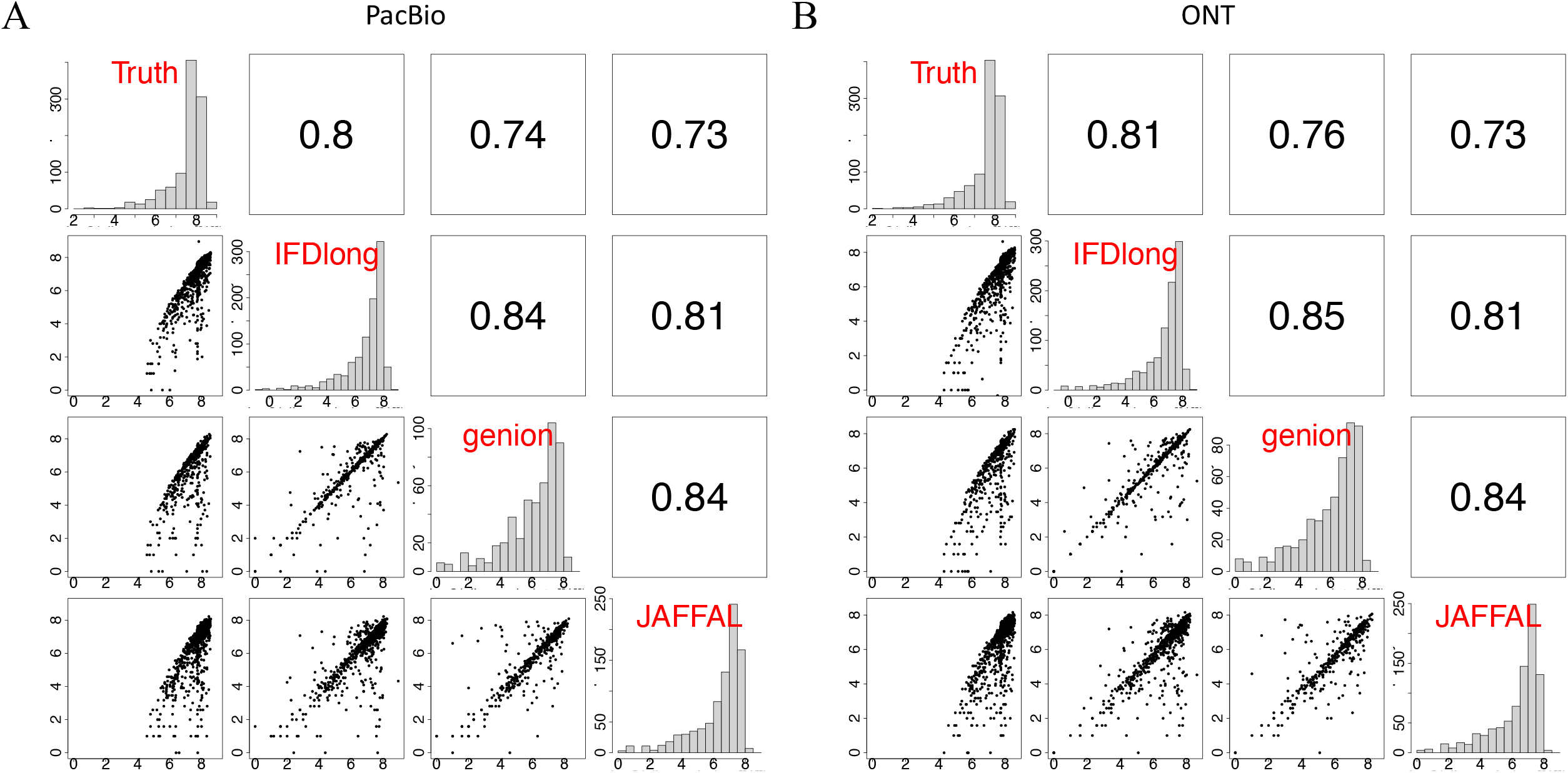
Pairwise comparison of fusion quantification in the Type F1 simulation. (A) The Type F1 simulation with median length of 1500 bp and PacBio accuracy. (B) The Type F1 simulation with median length of 1500 bp and ONT accuracy. The bottom left cells present the pairwise scatter plot of isoform expression. The upper right cells indicate the Spearman’s correlation.

**Figure S14:**
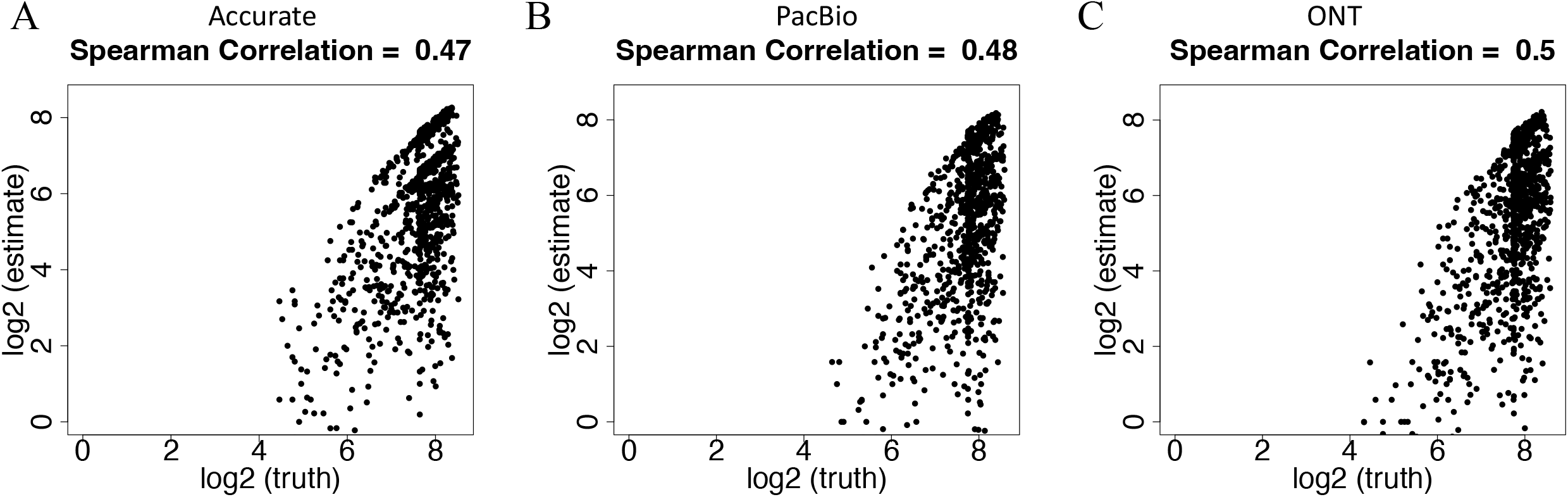
Scatter plot of isoform-level fusion quantification comparing the truth the IFDlong estimation in the Type F1 simulation. (A) The Type F1 simulation with median length of 1500 bp and accurate setting. (B) The Type F1 simulation with median length of 1500 bp and PacBio accuracy. (C) The Type F1 simulation with median length of 1500 bp and ONT accuracy.

**Figure S15:**
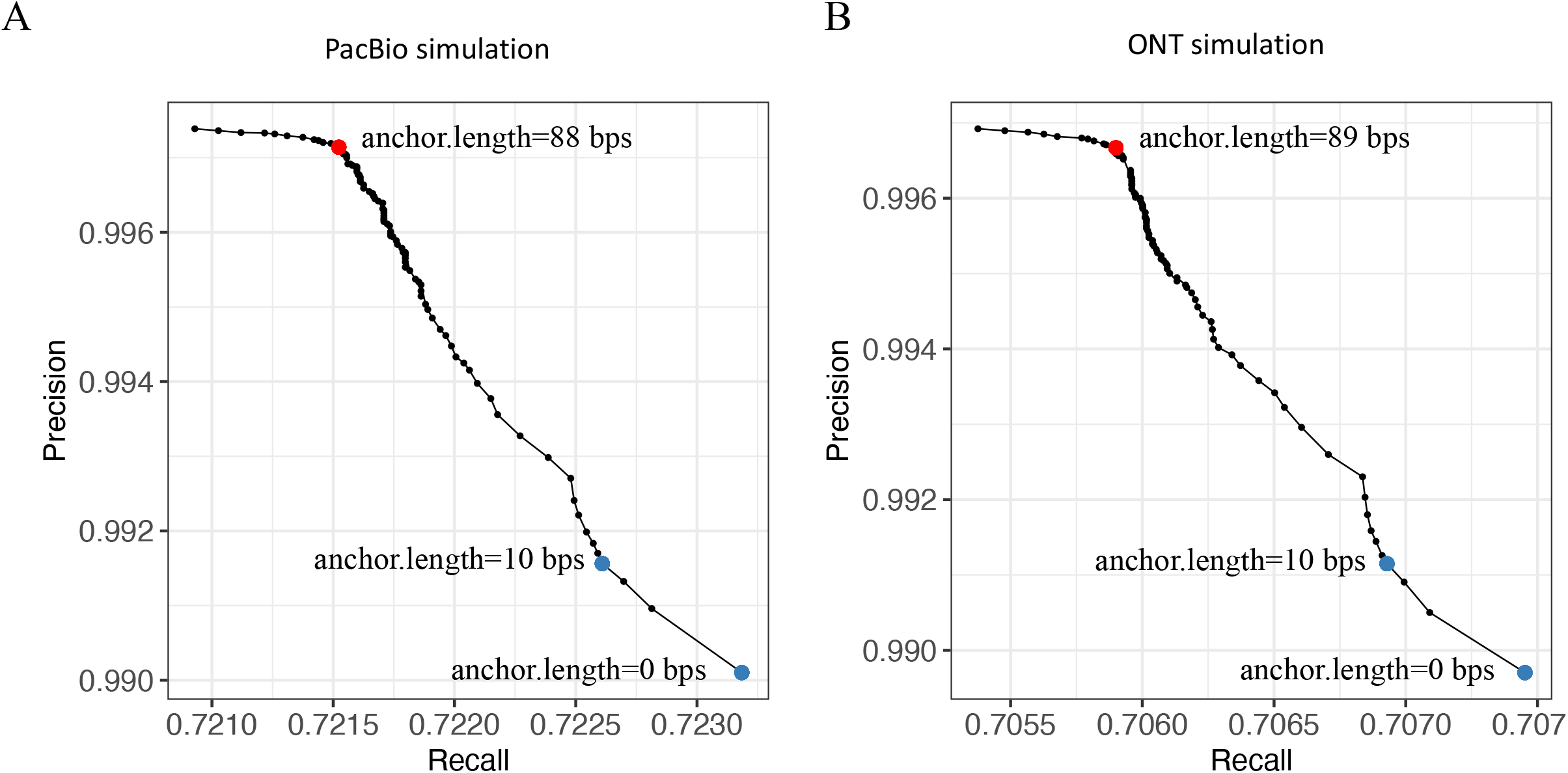
Precision and Recall curve for fusion transcript detection by IFDlong pipeline using different anchor lengths. (A) The Type F1 simulation with median length of 1500 bp and PacBio accuracy. (B) The Type F1 simulation with median length of 1500 bp and ONT accuracy.

**Figure S16:**
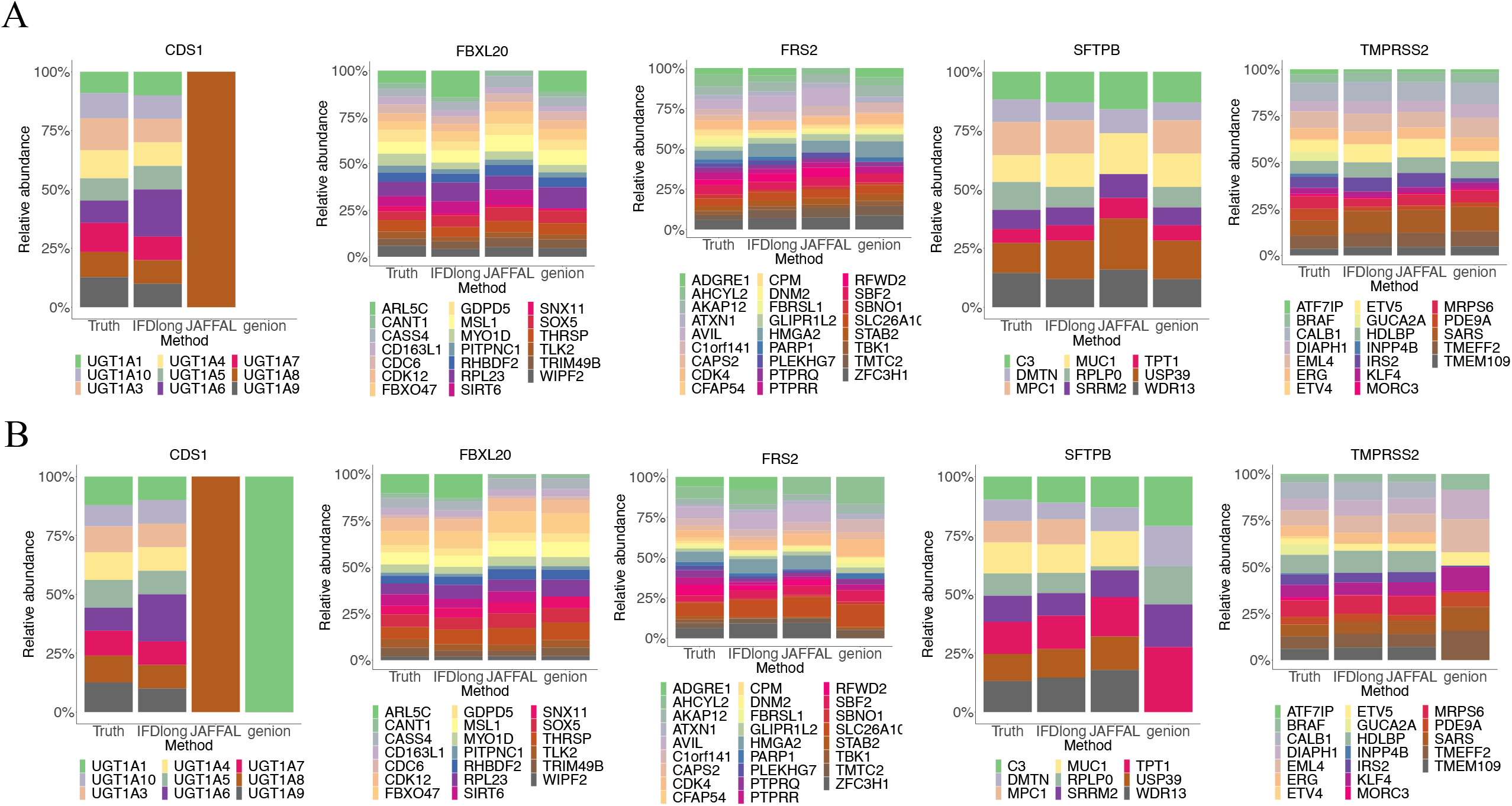
Relative abundance of multiple paired genes in the Type F2 simulation data with different accuracy setting. (A) The Type F2 simulation with 1500 bp median length and PacBio setting. (B) The Type F2 simulation with 1500 bp median length and ONT setting.

**Figure S17:**
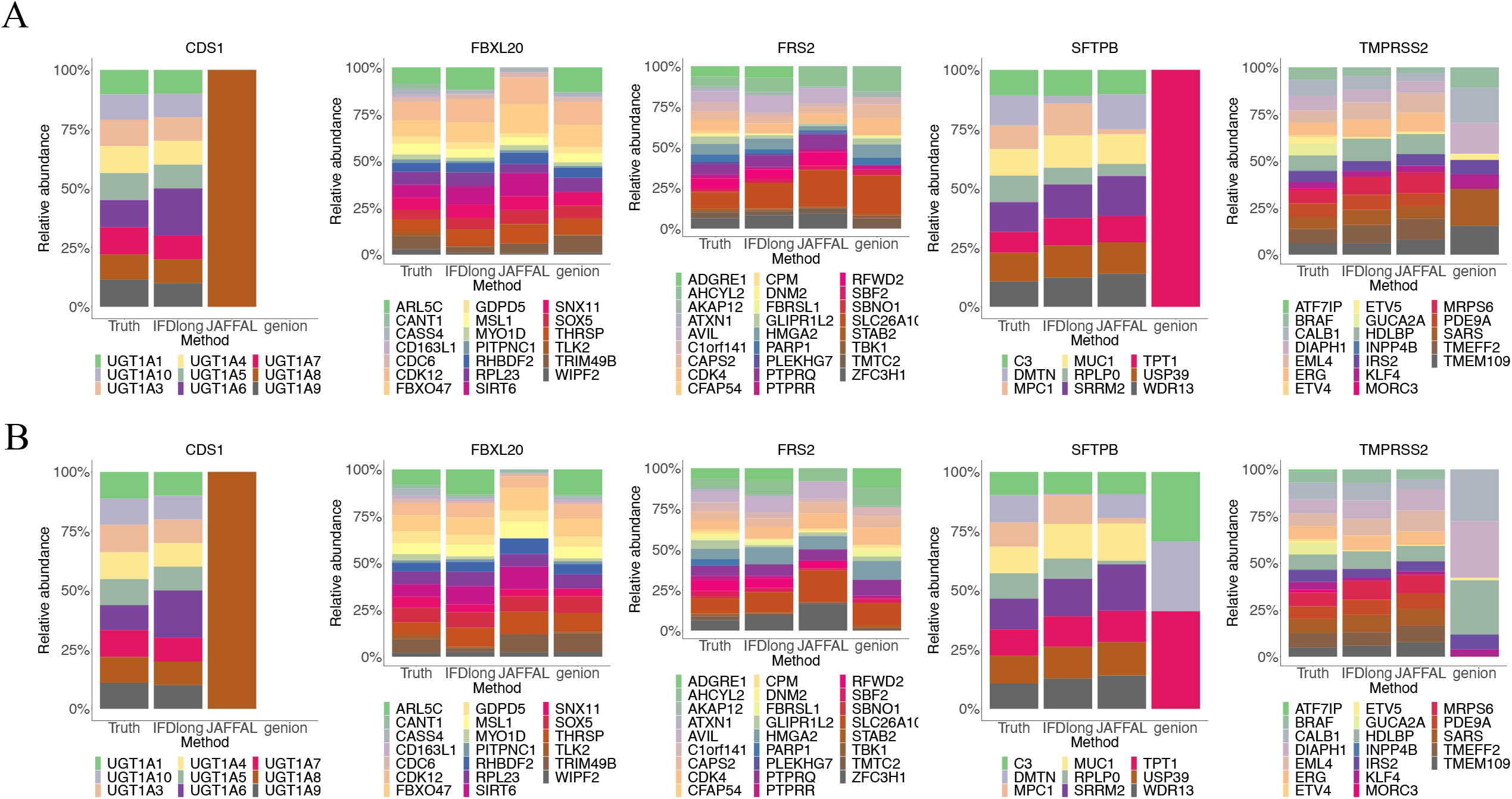
Cosine similarity of the relative abundance of multiple paired genes in the Type F2 simulation data. (A) The Type F2 simulation with median length to be 200bp and accurate setting. (B). The Type F2 simulation with median length to be 1000bp and accurate setting. (C) The Type F2 simulation with median length to be 1500bp and PacBio setting. (D) The Type F2 simulation with median length to be 1500bp and ONT setting.

**Figure S18:**
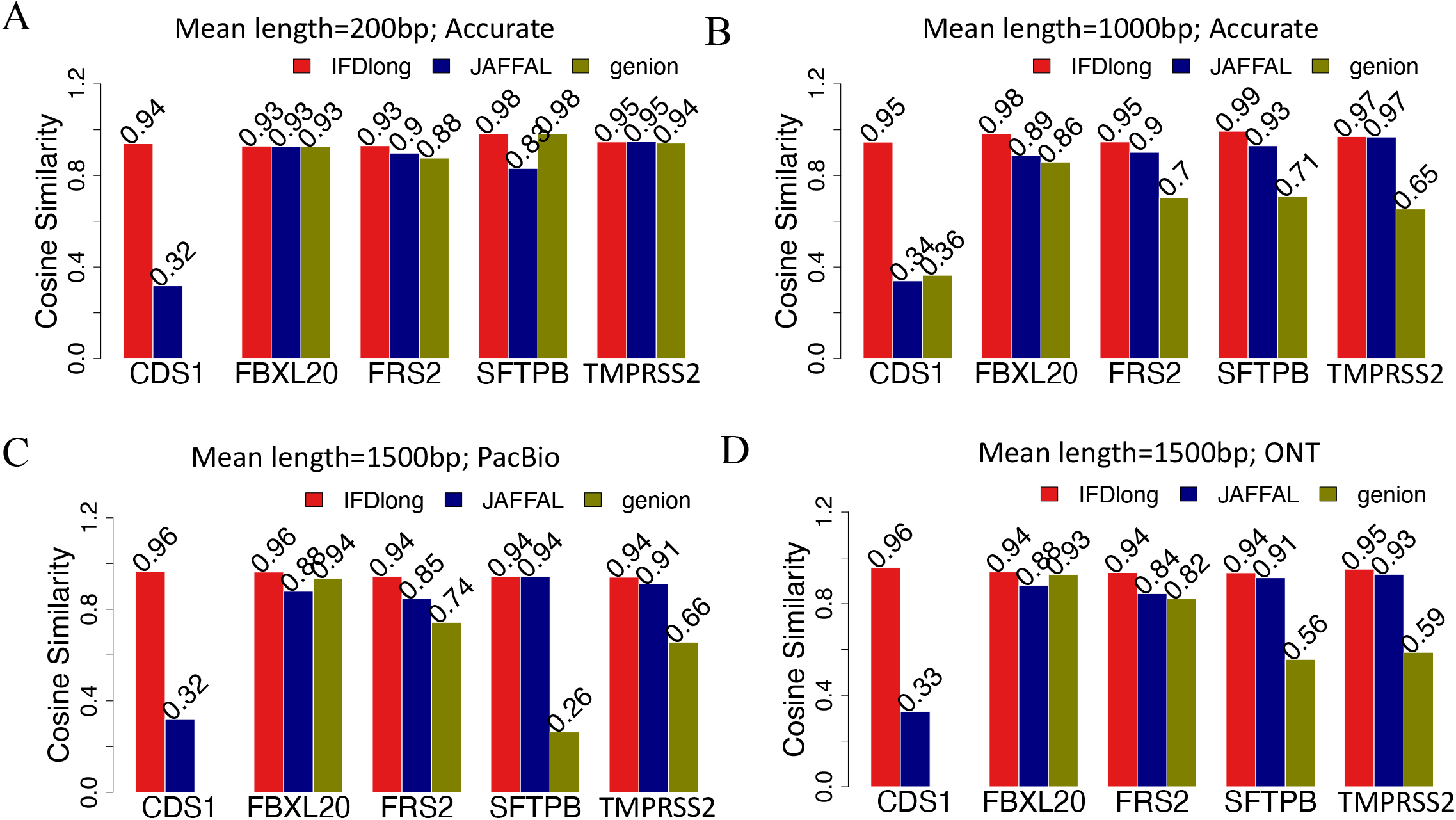
Relative abundance of multiple paired genes in the Type F2 simulation data with gradient median length. (A) The Type F2 simulation with 200 bp median length and accurate setting. (B) The Type F2 simulation with 1000 bp median length and accurate setting.

## Supplementary Table

**Table S1: Isoform quantification by IFDlong on UHR dataset**.

**Table S2: Isoform quantification by IFDlong on MCF7 dataset**.

**Table S3: Isoform quantification by IFDlong on mouse C2C12 dataset**.

**Table S4: Isoform quantification by IFDlong on mouse heart dataset**.

**Table S5: Differentially expressed isoforms in HCC dataset comparing tumor and normal. Table S6: Pathway analysis on the differentially expressed isoforms in HCC dataset**.

**Table S7: Popular fusion transcripts in TCGA dataset that were employed in the Type F2 simulation**.

**Table S8: Two-way fusions detected in MCF7 ONT dataset by IFDlong**.

## Notes

### Competing Interest Statement

The authors have declared no competing interest.

